# P2K1 P2K2 are required for early, mechanical adaptation to osmotic and cell wall stress in Arabidopsis roots

**DOI:** 10.64898/2026.10.15.759352

**Authors:** Dhika Amanda, Óscar A. Badias, Luis Alonso Baez, Maarten Besten, Joris Sprakel, Thorsten Hamann

## Abstract

The cell wall and plasma membrane (PM) form the interface between a plant cell and its environment yet how their mechanical properties change in response to stress is poorly understood. Here we investigate how the mechanical properties of the cell wall, plasma membrane, and vacuole change in vivo in *Arabidopsis thaliana* roots in response to hyperosmotic stress and cell wall damage (CWD) using a solvatochromic dye, molecular rotors, and Brillouin microscopy. We observe initially dynamic changes in mechanical properties of the PM that are sensitive to both hyperosmotic stress and CWD, while the cell wall changes are limited and the vacuole adapts only after several hours. In the *p2k1 p2k2* double mutant, lacking receptors for extracellular ATP perception, sorbitol-induced effects on PM mechanostasis and early isoxaben-induced increases in PM order are lost. Our results indicate that P2K1 and P2K2 are required to modulate adaptive changes in mechanical properties of the PM induced by hyperosmotic stress and CWD.

## Introduction

Plant cells are surrounded by a rigid yet dynamic cell wall that provides mechanical strength, controls cell shape, and contains the turgor pressure generated by osmotic forces. Turgor exerts an outward force on the plasma membrane (PM), increasing membrane tension ^1,2^. Changes in tension can reorganize lipids and proteins within the PM and modify PM fluidity impairing PM functionality. Membrane tension can rise further during hypoosmotic stress or as a result of cell wall damage (CWD) occurring during growth, development, or exposure to abiotic or biotic stress^3^. Conversely, low membrane tension can arise under conditions such as hyperosmotic stress and also impair PM functionality and nutrient transport, induce stress responses, and alter PM organization ^4–7^. Such reorganization involves changes in lipid synthesis and degradation, as well as modifications to membrane fluidity, to enable adaptation to diverse environmental conditions ^8–10^. Lipid remodelling frequently involves changes in fatty acyl chain length and degree of saturation ^6^. In Arabidopsis leaves, two major sphingolipid types—glucosylceramides (GlcCers) and glycosylinositol phosphorylceramides (GIPCs)—make up approximately 60% and 30% of total plasma membrane sphingolipids, respectively and were originally suggested to localize to lipid rafts ^11–13^. Recent work suggests that PM-localized GIPCs are crosslinked via boron (and Hechtian strands) with the cell wall, forming physical PM-cell wall attachments ^14^. Knowledge regarding plant PM organization has increased profoundly recently and standardized terminology has been agreed upon ^15^. Nanodomains have been defined as the smallest, discernible entity of PM organization above the level of single molecules ^15^. The contributions of nanodomains to different cellular functions and signalling activities during development and interactions with the environment have been reviewed recently ^6^.

Interest in the cell wall–PM continuum has increased recently since it forms the space where mechanical stimuli, osmotic imbalance, changes in PM tension, and cell wall integrity impairment are detected ^16–19^. Cellulose synthase complexes, together with cellulose microfibrils and Hechtian strands, appear to play a major role in coordinating interactions between the cell wall and PM during hyperosmotic stress^20^. Hechtian strands form when plant cells are exposed to hyperosmotic stress and the PM begins to detach from the cell wall. Their formation can result in increased local membrane tension near the adhesion site ^21,22^ while overall membrane tension may decrease ^23^. Recent work showed that overexpression of LecRK-I.9/P2K1 increases Hechtian strand formation, whereas loss of P2K1 function has limited phenotypic effects ^24^. The authors proposed that P2K1 acts as “a molecular glue” helping to anchor the PM to the cell wall ^20^. This concept is supported by previous evidence that implicated P2K1 originally in PM–cell wall adhesion and RGD-peptide perception, thereby linking it to cell wall integrity and mechanosensing ^25,26^. However, substantial experimental evidence has accumulated implicating P2K1— together with P2K2/LecRK-I.5—in perception of extracellular ATP (eATP), which is released during plant exposure to biotic or abiotic stress and acts as damage associated molecular pattern ^27–31^. Importantly, only simultaneous loss of both P2K1 and P2K2 abolishes most of the eATP-induced responses^29^.

Several new technologies have been developed recently that allow in vivo characterization of mechanical properties of cells with subcellular resolution ^32,33^. Amongst them are molecular rotors of the boron-dipyrromethene (BODIPY/BDP) type, which are targeted to specific sub-cellular structures ^34,35^. Molecular rotors are rigidochromic fluorescent dyes that undergo intramolecular rotation upon photoexcitation. The rotation rate depends on the physical constraints of the surrounding environment, making both fluorescence lifetime and intensity sensitive to local micromechanical properties, or more specifically to the local molecular free volume ^36,37^. BODIPY probes targeted to the vacuole (VC: sulfo-BDP) enable us to analyse the viscosity (i.e. degree of crowding) inside of the vacuole. Increased lifetime means increased viscosity in the vacuole while decreases mean reduced viscosity. Increased lifetime in the rotor targeted to the PM (N+-BDP) indicates reduced PM fluidity while decreased lifetime indicates increased fluidity. Importantly, PM fluidity as reported by this probe also provides indirectly insights into PM tension because the lifetime of the fluorescent molecular rotor increases with higher lipid packing density, indicating a greater proportion of Liquid-ordered (Lo) lipid and reduced membrane tension. On the contrary, a decreased lifetime indicates lower lipid packing density, a higher proportion of Liquid disordered (Ld) lipid, and increased membrane tension. The cell wall-targeted CarboTag-BDP rotor measures the nanometric porosity in the cell wall mesh, ie. how much free volume is available for the probe to undergo its rotation ^34^. Reduced porosity leads to increased lifetime, while increased porosity leads to decreased lifetime. Rotation seems to be mainly determined by the level and crosslinking state of pectins, to which the probe binds mainly. C-Laurdan is another probe used to investigate the lateral organization of the PM^38–40^. It is a solvatochromic probe, which displays significant changes in its dipole moment upon excitation and is sensitive to the presence and mobility of water dipoles within a lipid bilayer and its surrounding two aromatic rings, causing a shift in the emission spectrum. Since Ld lipids permit more water molecules to enter the bilayer (polar), while Lo lipids restrict water entry (non-polar), the C-Laurdan emission spectrum shifts in a corresponding manner ^41,42^. Accordingly, shifts in the spectrum of C-Laurdan provide insights into the lateral organization of the PM, which can be quantified using the Generalized Polarization (GP) value. The GP value quantifies the ratio of spectral shift of fluorescence emission at two distinct wavelengths (440 nm and 490 nm) with higher GP values indicating a more ordered membrane and lower GP values indicating a more fluid membrane ^43^.

A complementary method to explore changes to plant micromechanical properties is Brillouin microscopy ^32,44,45^. This non-invasive, label-free technology quantifies changes in linewidth and detects frequency shifts between the elastic and inelastic scattered light in samples. Brillouin microscopy enables this analysis by detecting spontaneous thermally induced density fluctuations in the samples investigated. This is possible because the fluctuations induce local volume changes by transiently compressing the material. The Brillouin frequency shift is proportional to the longitudinal modulus, which reports on how strongly the material resists the induced volume changes, while the linewidth is proportional to the longitudinal viscosity, which indicates how quickly fluctuations lose coherence, reflecting viscous processes ^32,44,45^. Changes in frequency shift are indicative of modified stiffness while linewidth changes are providing insights into viscosity. By combining the three different technologies, we can investigate in vivo PM lateral organization (C- Laurdan), PM fluidity and tension (N+-BDP), vacuolar cytoplasm viscosity (sulfo-BDP), cell wall porosity (CarboTag-BDP), and longitudinal modulus/viscosity at the cell borders and the cytoplasm (Brillouin microscopy). This provides a comprehensive picture of the micromechanical state of the tissue.

Here we apply hyperosmotic stress (induced by sorbitol) and CWD (induced by isoxaben) to investigate how different types of stress affect the mechanical properties of the cell wall, PM, cytoplasm and vacuole in vivo. We determine in *Arabidopsis thaliana* seedling roots how PM fluidity changes in different cell types and developmental stages upon exposure to hyperosmotic stress and CWD. We perform time-course experiments focusing on cortex cells and determine how mechanical properties of cell walls, PM, and vacuole change in response to hyperosmotic stress and CWD. Lastly, we investigate how loss of the P2K1 P2K2 receptor kinases, implicated in eATP perception and mediating adaptive changes in the cell wall and PM, modulate mechanical properties of different subcellular structures and affect changes induced by the different stress types.

## Results

### Hyperosmotic stress induces adaptive changes in mechanical properties of sub-cellular structures

Previous studies have examined how hyperosmotic stress affects plasma membrane organization within specific cell types or developmental stages. Here, we set out to investigate the impact of hyperosmotic stress on plasma membrane organization across multiple tissue layers and developmental stages simultaneously, using the *Arabidopsis thaliana* seedling root as a model system. For this purpose, we used the PM-localized molecular rotor N+-BDP ^35^ to generate PM fluidity maps of seedling root tips that were either mock-or sorbitol (0.3M)-treated for three hours (Fig. 1). We have previously shown that these experimental conditions induce a transient increase in abscisic acid production, peaking at three hours, which is a well-established response to hyperosmotic stress ^46^.

**Figure 1.**
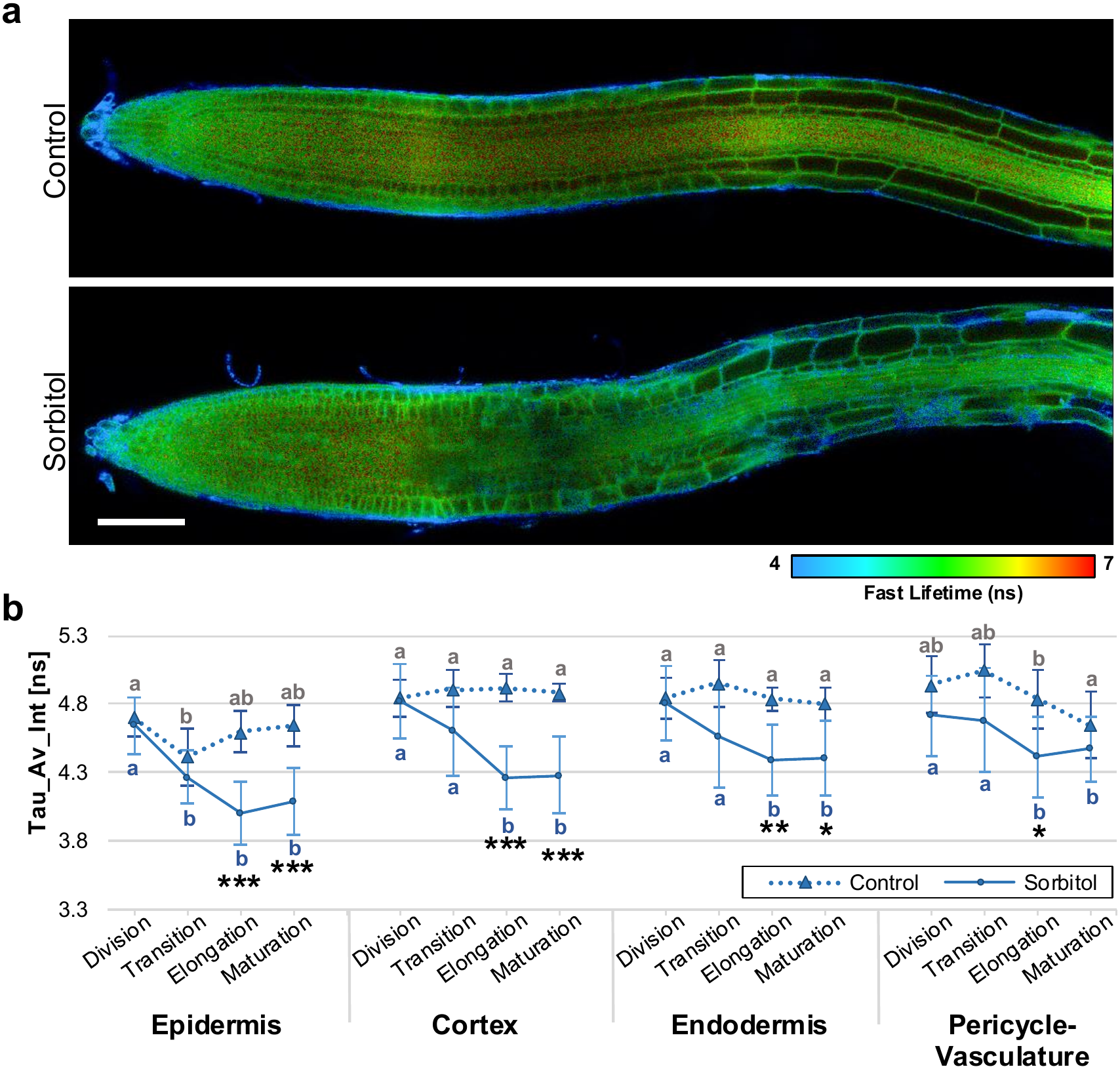
Plasma membrane fluidity in different cell layers of *Arabidopsis thaliana Col-0* seedling roots exposed to hyperosmotic stress. **a**, N+-BDP-based fluorescence lifetime map of the plasma membranes in Col-0 seedling roots mock- or 0.3M sorbitol-treated for three hours. Scale bar = 100μm. **b,** Mean fluorescence lifetime of N+-BDP plasma membranes of Col-0 seedling roots after mock or 0.3M sorbitol treatment. Error bar = standard deviation. Statistics: one-way ANOVA with Post Hoc – Tukey HSD. X axis indicates different developmental zones for each cell layer. Y axis indicates average fluorescence lifetime (in nanoseconds). *Asterisks* indicate significant difference between mock or sorbitol treated roots in each individual zone \**P*<0.01, \*\**P*<0.003, \*\*\**P*<0.001; *letters* indicate significant differences between zones within the same cell type and treatment group. n = 10, except in mock division and transition zones n = 9.

Mock- and sorbitol-treated root tips displayed differences in cell shape, which were most pronounced in the epidermis (Fig. 1a). We compared PM fluidity both within individual tissue layers (i.e., along the developmental axis) and between mock- and sorbitol-treated samples within a given developmental stage (Fig. 1b).

Comparisons between tissue layers revealed differences between the epidermis, cortex, and endodermis in the elongation and maturation zones, whereas differences in the pericycle-vasculature layer were less pronounced. Within-layer comparisons showed that the epidermis already differed in the transition zone, while the cortex, endodermis, and pericycle-vasculature layers showed differences only in the elongation and maturation zones. Together, these results indicate that the PMs of all cell layers examined respond to hyperosmotic stress, and that the extent of the response may be influenced by the developmental stage.

Having established that hyperosmotic stress affects PM fluidity similarly across cell layers, we next asked whether cell shape influences mechanical properties, using BODIPY-based molecular rotors (Supplementary Fig. 1 and 2) decorated with distinct chemical groups to target different structures. We focused on the cortex as a representative cell layer and used three molecular rotors targeting the vacuole (sulfo-BDP), the plasma membrane (N+-BDP), and the cell wall (CarboTag-BDP). To distinguish between two cortical cell shapes in the early and late elongation zones, we calculated the ratio of longitudinal to transversal cell wall length, classifying cells as quadratic (ratio of 1–1.5) or rectangular (ratio > 1.5).

Although minor differences were observed, the cell wall, plasma membrane, and vacuole showed similar responses in both quadratic and rectangular cells. We therefore combined the datasets for subsequent analyses (Fig. 2 and Supplementary Fig. 2).

**Figure 2.**
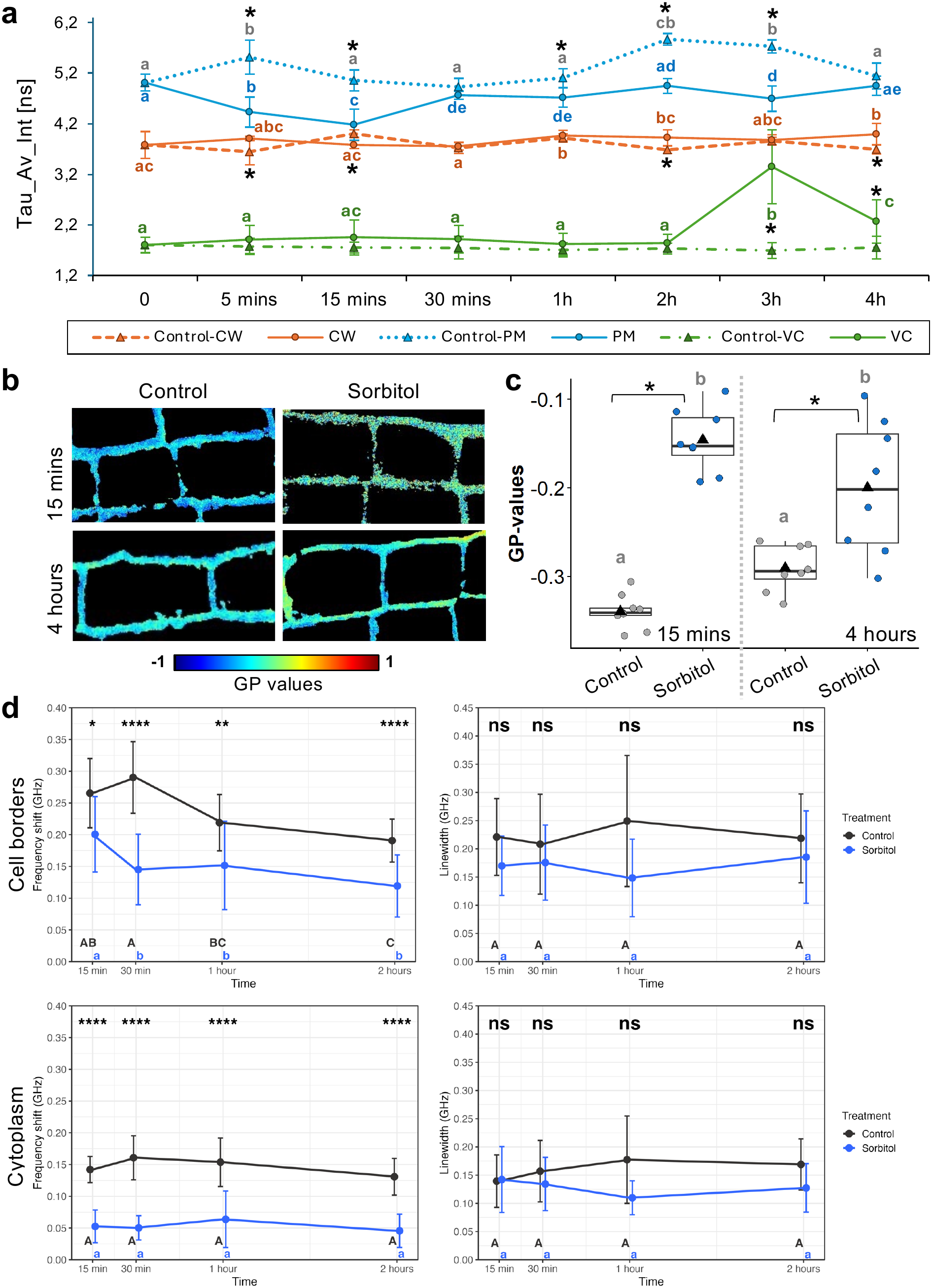
Physicochemical properties of subcellular structures in Col-0 cortex cells exposed to hyperosmotic stress. **a**, Mean fluorescence lifetimes of sulfo-BDP (VC: vacuole), N+-BDP (PM: plasma membrane) and CarboTag-BDP (CW: cell wall) in mock- or 0.3M sorbitol-treated cortex cells. X axis indicates treatment periods, Y axis indicates average fluorescence lifetime. From left to right, respectively, control- PM n = 10, 8, 10, 8, 9, 8, 10, 8; all treated-PM n = 10; all control-CW n = 10, except 3h n = 9; treated-CW n = 10, except control 4h = 8; all VC n = 10, except control 4h n = 8 and sorbitol 4h n = 9. **b,** Graphical representation of PM polarity of mock- or 0.3M sorbitol-treated cortex cells (magnified from Supplementary Fig. 8) stained with C-Laurdan. Generalized polarisation (GP) values are color-coded and range from -1 to 1. **c,** GP values of C-Laurdan-stained cortex cell PMs (n = 9 roots, except control 15 mins n = 8). **d,** Frequency shift and linewidth of cytoplasm and cell borders of mock or sorbitol-treated cortex cells detected with Brillouin light scattering microscopy (controls n = 9, except 1h n = 10 and 2h n = 14 roots. 15 mins – 2h sorbitol n = 16, 17, 21, and 19 roots). Error bars = standard deviation. *Asterisks* indicate statistical significance of differences between mock and sorbitol \**P*<0.001, **P<0.003, ***P<0.01, ****P<0.0001; *letters* indicate significant differences between time points of same treatment type. Statistics a-c: one-way ANOVA with Post Hoc – Tukey HSD.

Next, we performed time-course experiments, comparing mock- and sorbitol-treated samples at each individual time point (indicated by stars, with corresponding p-values) as well as changes over time within each treatment for a given cell layer (indicated by letters). PM fluidity increased transiently after five and fifteen minutes and fluctuated over time possibly due to fresh medium effects. In response to hyperosmotic stress, fluidity decreased after five and fifteen minutes, recovered (relative to timepoint 0) and remained stable for the remainder of the treatment period (Fig. 2a). Cell wall porosity (CarboTag-BDP) exhibited minor changes over time which were significantly different at certain timepoints. Vacuolar cytoplasm viscosity increased transiently after three hours, implying an increase in crowding.

To complement the PM fluidity results generated with BODIPY, we investigated the lateral organization, i.e. chemical polarity, of the PM using the polarity probe C-Laurdan at early (15 minutes) and late (4 hours) timepoints (Fig. 2b, c). The time points were chosen since they seem to represent early and late responses to the stress. Exposure to sorbitol for both 15 minutes and 4 hours reduced PM polarity, as shown by the polarity map (Fig. 2b) and by elevated GP values relative to mock (Fig. 2c). This effect appeared more pronounced after 15 minutes than after 4 hours, suggesting that the lateral organization of the PM adapts to hyperosmotic stress over time.

We next performed time-course experiments with mock- or sorbitol-treated *Arabidopsis* seedling root tips, using Brillouin microscopy to examine how frequency shift and linewidth change at cell borders and in the cytoplasm of cortex cells (Fig. 2d). Linewidth did not differ substantially in the cytoplasm or at the cell border between mock- and sorbitol-treated cortex cells. Frequency shift, however, differed between mock- and sorbitol-treated cells as early as 15 minutes similar to effects observed for PM fluidity ie. sorbitol-and mock-treated cells showed initially opposite patterns. The differences persisted over time. This suggests that mock and sorbitol treatments have rapid, qualitatively different effects on cell borders. To summarize, results from three different methodologies consistently detect changes in the mechanical properties of cortex cell borders within minutes of exposure to hyperosmotic stress. Changes in vacuolar viscosity occur hours later, highlighting that different subcellular structures respond to hyperosmotic stress on different timescales.

### CWD and hyperosmotic stress have distinct effects on mechanical properties of subcellular structures

Our results indicate that hyperosmotic stress induces dynamic alterations in the mechanical properties of cortex cells. Recent work indicated that CWD, caused by the cellulose synthase inhibitor isoxaben (ISX), also affects PM organization and cellulose synthase association with the PM ^47–50^. We have shown before that co-treatments with osmotica suppress most of the CWD-induced responses ^51^. We therefore investigated using the same experimental approaches as before, how ISX-induced CWD and combined treatments of ISX/sorbitol affect mechanical properties of Arabidopsis seedling root tips.

We examined the effect of ISX and ISX/sorbitol treatments on PM fluidity across whole root tips (Fig. 3, Supplementary Figs. 3–4) and cortex cells specifically (Fig. 4). Whole root tip (Fig. 3) and cortex time-course (Fig. 4) measurements use are affected by the focal plane selected, use different ROI definitions and are analysed separately (see materials and methods for details), making direct comparisons between results presented in figures 3 and 4 difficult. Because ISX is dissolved in DMSO, the mock control also contained DMSO. DMSO-treated root tips showed no pronounced differences between cell layers, though PM fluidity appeared reduced at later developmental stages (Fig. 3b). ISX-treated root tips showed lower PM fluidity than DMSO or sorbitol-treated root tips across all cell layers and developmental stages (Fig. 3b). While the sorbitol-treated transition zone did not differ from mock, the ISX/sorbitol-treated transition zone differed from the DMSO control in all examined cell layers. Next, we performed time-course experiments to examine how ISX and ISX/sorbitol treatments affect the mechanical properties of cortex cells (Fig. 4a, b). Unlike sorbitol-treated cortex cells, which showed an early, transient increase in PM fluidity, ISX- and ISX/sorbitol-treated cells did not (compare Fig. 4a, b with Fig. 2a). However, PM fluidity did increase at later timepoints in response to ISX and ISX/sorbitol treatment, in contrast to sorbitol alone. Cell wall porosity increased transiently between one and three hours of ISX treatment, but this transient increase was absent in cortex cells co-treated with ISX/sorbitol. Vacuolar viscosity appeared to increase earlier than in sorbitol-treated cortex cells (compare Fig. 4a, b with Fig. 2a). Together, these observations suggest that ISX and sorbitol have distinct, time-dependent effects on the micromechanical features of subcellular structures. Because combined treatments do not produce additive/contrary effects, ISX and sorbitol likely act on connected processes; notably, combining the treatments suppresses certain sorbitol-specific effects (namely, the early increase in PM fluidity). Vacuolar viscosity again appeared to change last, suggesting that the responses occur in a sequential manner.

**Figure 3.**
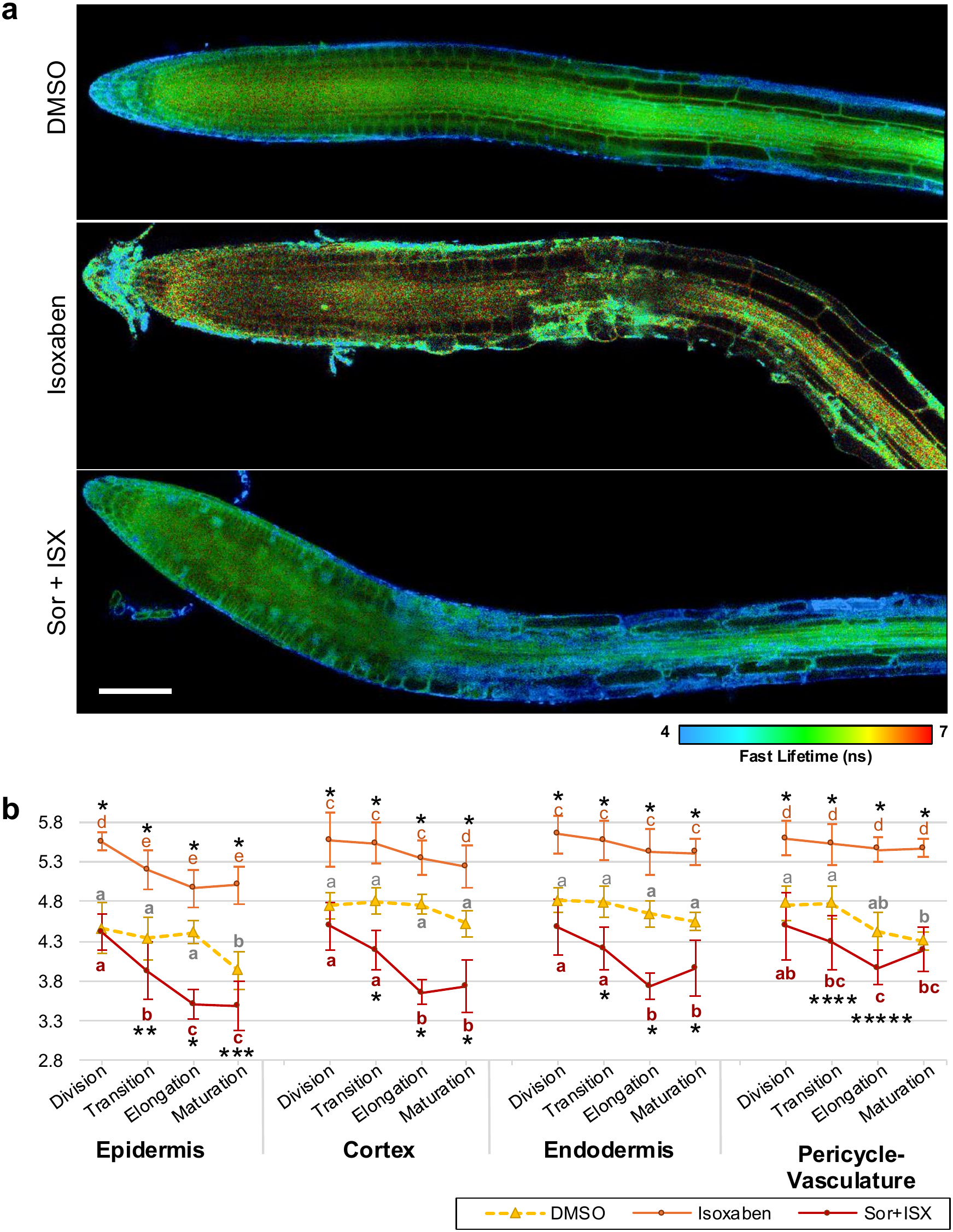
Plasma membrane fluidity in different cell layers of Col-0 seedling roots exposed to different stress types. **a**, N+-BDP-based fluorescence lifetime map of the PMs in Col-0 seedling roots mock (DMSO), 0.6μM isoxaben or 0.3M sorbitol/0.6μM ISX-treated for three hours. Scale bar = 100μm. **b,** Mean fluorescence lifetime of N+-BDP in PMs of Col-0 seedling roots after mock, 0.6μM isoxaben or 0.3M sorbitol/0.6μM ISX treatments. Y axis indicates average fluorescence lifetime (in nanoseconds). X axis indicates different developmental zones for each cell layer. Error bars = standard deviation. Statistics: one-way ANOVA with Post Hoc – Tukey HSD. *Asterisks* indicate significant difference between mock and ISX- or sorbitol + ISX-treated roots in each individual zone \**P*<0.001, \*\**P*=0.044, \*\*\**P*=0.021, \*\*\*\**P*=0.009, \*\*\*\*\**P*=0.024; *letters* indicate significant differences between zones within the same cell type and treatment group. Roots treated n = 10 for each treatment with exception of DMSO n = 9.

**Figure 4.**
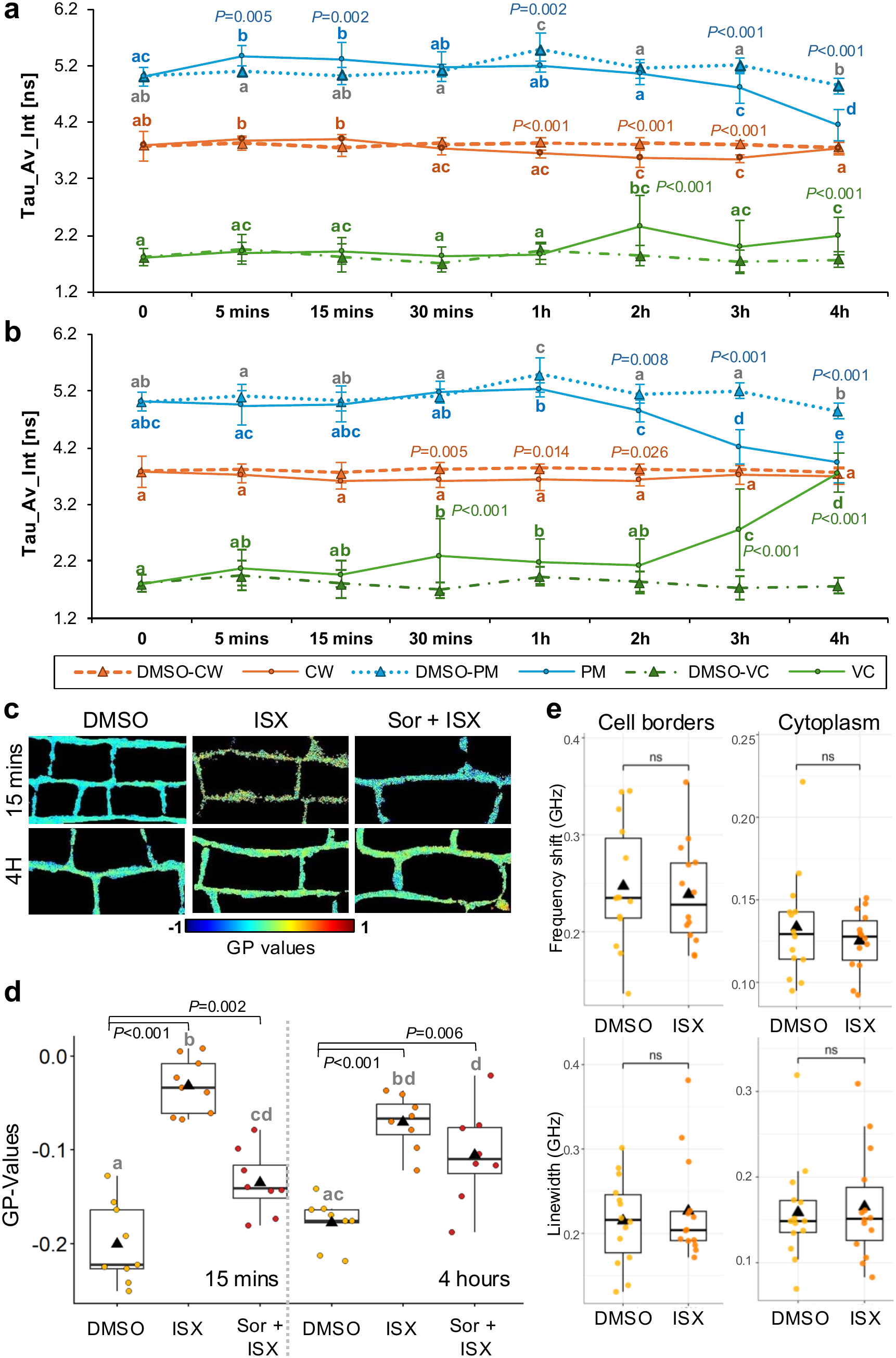
Physicochemical properties of subcellular structures of Col-0 cortex cells in the root elongation zone exposed to CWD and/or hyperosmotic stress. **a**, Mean fluorescence lifetimes of sulfo-BDP (VC), N+-BDP (PM) and CarboTag-BDP (CW) in mock (DMSO)- or 0.6μM ISX-treated cortex cells. From left to right, respectively, all DMSO-PM n = 10, except 1h n = 8; treated-PM n = 10, 8, 8, 10, 10, 10, 10, 8; all CW n = 10; DMSO-VC n = 10, 9, 10, 10, 8, 8, 8, 9; treated-VC n = 10, except 4h n = 8 roots. **b,** Mean fluorescence lifetimes of sulfo-BDP, N+-BDP, and CarboTag in mock (DMSO) - or 0.3M sorbitol + 0.6μM ISX- treated cortex cells. From left to right, respectively, treated-PM n = 10, 10, 10, 9, 9, 9, 10, 10; treated-CW n = 10, 10, 9, 10, 10, 10, 10, 11; treated-VC n = 10, except 30 mins n = 9 roots; mock (DMSO) is the same as in **a**. **c**, Graphical representations of cortex cells polarity (digitally magnified from Supplementary Fig. 8) after mock (DMSO), 0.6μM ISX, 0.3M sorbitol/0.6μM ISX treatments and stained with C-Laurdan. **d,** GP values of cortex cells PMs. Each data point indicates mean GP-values per individual root (all 15 mins n = 9, except Sor + ISX n = 8; 4 hours n = 8, except DMSO n = 9. *P*-values indicate statistical significance of differences between treated roots and their respective controls. *alphabet* refers to significance between all samples. **e,** Frequency shift and linewidth in cortex cells of seedling roots treated with DMSO or ISX for 15 minutes (n = 14 roots for each treatment, Statistics: Welch t-test). Statistics (**a**-**d**) were done using one-way ANOVA with Post Hoc – Tukey HSD. Error bars = standard deviation.

We next examined the lateral organization of cortex cell PMs in response to ISX and ISX/sorbitol treatments after 15 minutes and four hours with C-Laurdan (Fig. 6b). Comparing mock controls with and without DMSO, DMSO alone appeared to make the PM slightly less polar (Fig. 6b). Adding ISX increased the GP value after 15 minutes, indicating a further reduction in PM polarity. In ISX/sorbitol-treated cortex cells, the GP value fell between that of DMSO- and ISX-treated samples, suggesting that sorbitol suppresses the effect of ISX on PM polarity. After four hours, the ISX effect remained detectable, but the apparent suppressive effect of sorbitol was less pronounced. Brillouin microscopy analysis of cortex cells treated for 15 minutes detected no changes in frequency shift or linewidth at cell borders or in the cytoplasm (Fig. 4e). In previous work, we investigated frequency shift and linewidth under the same experimental conditions and detected a change in frequency shift in cortex cytoplasm and the cortex-endodermis border after three hours of ISX treatment ^52^. Together, these results suggest that ISX and sorbitol affect PM lateral organization in different ways: sorbitol appears to reduce the effect of ISX on PM organization, while ISX appears to suppress the early effect of sorbitol on fluidity. Unlike sorbitol, ISX does not induce frequency shifts (i.e., stiffness changes) in the cytoplasm or at cell borders at the early time point examined here.

### LecRK-I.9/P2K1 LecRK-I.5/P2K2 modulate PM fluidity and lateral organization

LecRK-I.9/P2K1 has been implicated in perceiving eATP and modulate the PM connection with the cell wall in *Arabidopsis* through Hechtian strands ^26,30^. Previous work suggested that LecRK-I.9/P2K1 and LecRK-I.5/P2K2 are both required for eATP perception and that only in *p2k1 p2k2* plants eATP perception is nearly abolished ^29^. Therefore, we decided to elucidate the role of LecRK-I.9/P2K1 and LecRK-I.5/P2K2 in regulating the mechanical properties of cortex cells exposed to sorbitol and ISX by characterizing *p2k1 p2k2* seedlings. We used the same experimental approaches as before, focusing on representative early (15 minutes) and late (four hours) time points (Fig. 5 and Supplementary Figs. 5-9). We used CarboTaq-BDP to characterize *p2k1 p2k2* cell walls and detected slightly increased porosity compared to the Col-0 under mock conditions (Fig. 5a). Both exhibited similarly increased porosity after 15 minutes of hyperosmotic stress (Fig. 5a). After four hours, porosity was similar in mock-treated Col-0 and *p2k1 p2k2* cell walls while in response to sorbitol only Col-0 cell wall porosity changed (Supplementary Fig. 7a). The *p2k1 p2k2* PM exhibited reduced fluidity compared to Col-0 in mock conditions after 15 minutes (Fig. 5a). However, no change in PM fluidity was detected after sorbitol treatment in *p2k1 p2k2* in contrast to Col-0 (Fig. 5a). After four hours, fluidity is similar in Col-0 and *p2k1 p2k2* cortex cells in both mock and sorbitol-treated PMs. Viscosity of Col-0 and *p2k1 p2k2* vacuoles is similar in mock and sorbitol conditions after 15 minutes (Fig. 5a). Viscosity was elevated in the sorbitol-treated *p2k1 p2k2* vacuole compared to Col-0 after four hours (Supplementary Fig. 7). After 15 minutes of DMSO mock treatment, the micromechanics of cell wall, PM and vacuole in *p2k1 p2k2* were altered compared to Col-0 (Fig. 5a). This suggests hypersensitivity of the *p2k1 p2k2* cortex cells to any challenge affecting lipid PM components. The micromechanical properties of the cell wall, vacuole or PM did not change in Col-0 in response to ISX. Cell wall porosity remained unchanged in *p2k1 p2k2* but vacuolar viscosity was decreased while PM fluidity was increased in response to ISX. After four hours of DMSO treatment, *p2k1 p2k2* vacuole viscosity and cell wall porosity were similar to Col-0, but PM fluidity was lower (Supplementary Fig. 7). In cell walls, no porosity changes were detected in ISX- or ISX/sorbitol-treated cortex cells. PM fluidity was increased both in Col-0 and *p2k1 p2k2* in response to ISX and ISX/sorbitol-treatments (Supplementary Fig. 7b). To summarize, the results suggest that P2K1 P2K2 are required for PM mechanostasis and early adaptive changes induced by hyperosmotic, DMSO, and ISX-derived stress. P2K1 P2K2 seem to be required for long term adaptation of cell wall and vacuole micromechanics to hyperosmotic stress. Vacuole viscosity exhibits delayed changes, suggesting it may adapt to earlier events taking place at the PM and cell wall.

**Figure 5.**
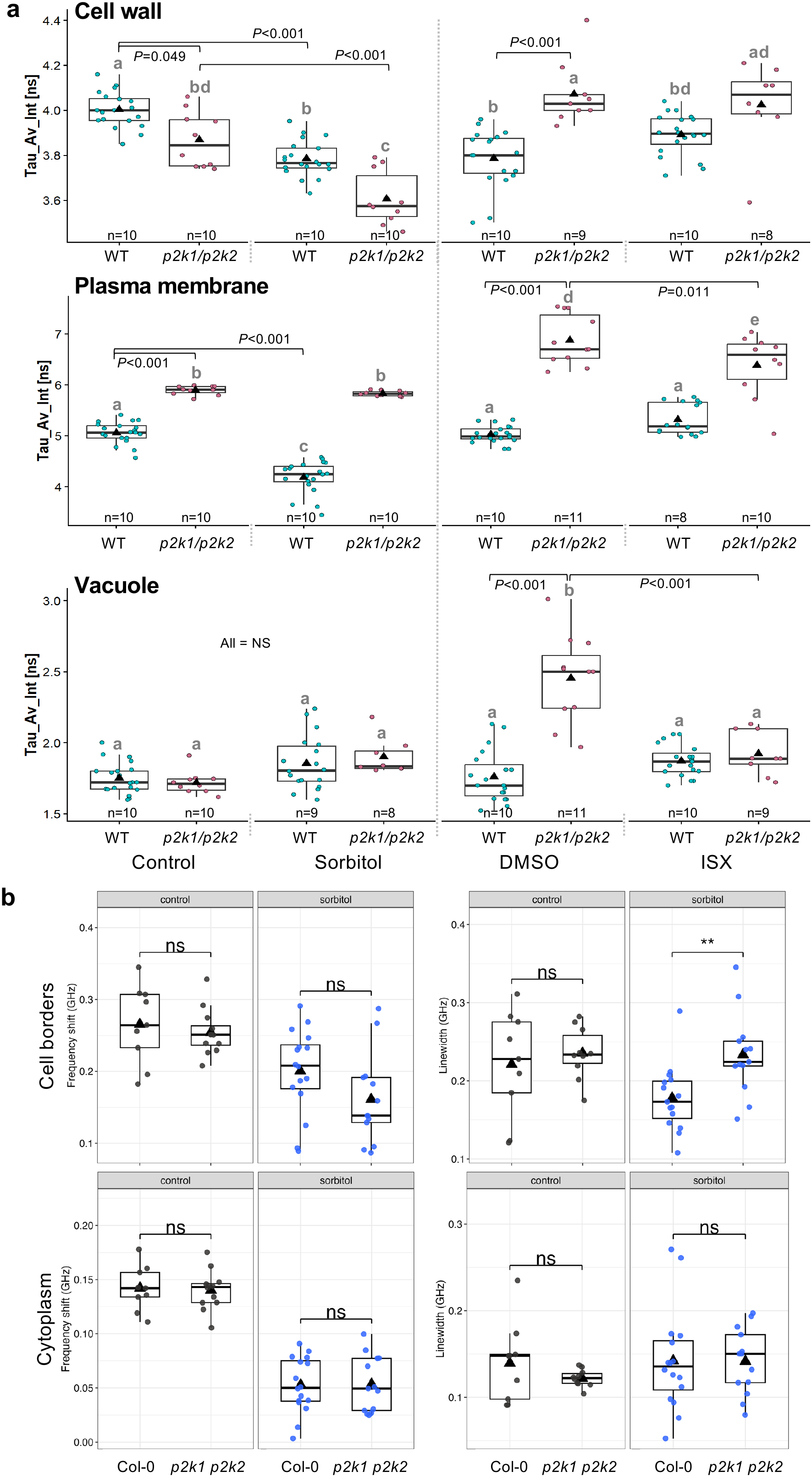
Mechanical properties of subcellular structures of Col-0 and *p2k1/p2k2* cortex cells in the root elongation zone. **a**, Mean fluorescence lifetimes of cell walls, PMs, and vacuoles in cortex cells treated with of 0.3M sorbitol and 0.6μM ISX for fifteen minutes compared to control and mock (DMSO). **b,** Brillouin light scattering microscopy-derived frequency shift and linewidth of cytoplasm and cell borders of mock or 0.3M sorbitol-treated cortex cells (control n:12, sorbitol n:13). Statistics in **a**: one-way ANOVA with Post Hoc – Tukey HSD, *P*-values indicate statistical significance of differences between treated roots and their respective controls, *alphabet* refers to significance between all samples. Statistics in **b** Welch’s t-test. Error bars = standard deviation.

**Figure 6.**
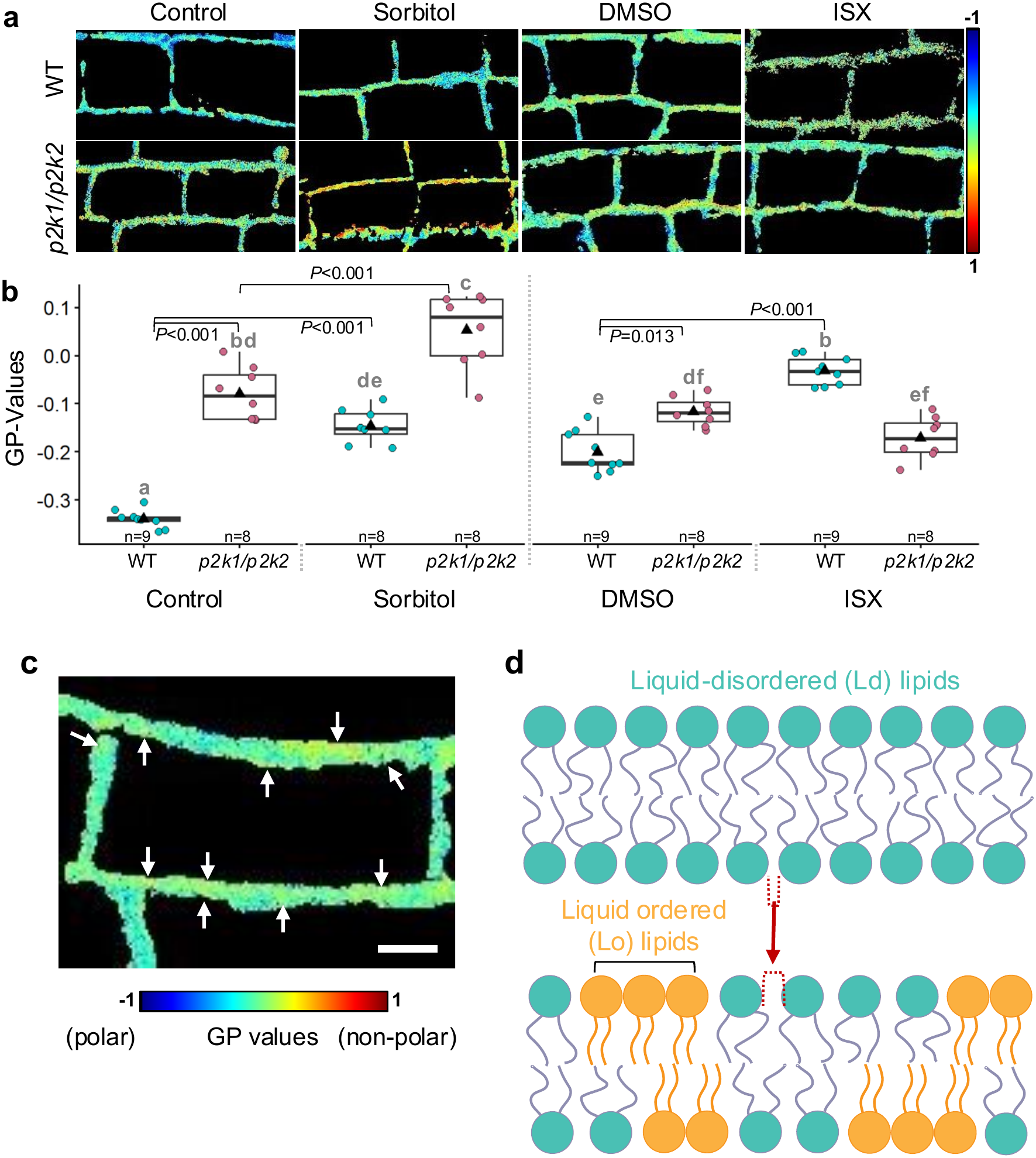
PM polarity in Col-0 and *p2k1/p2k2* cortex cells exposed to different stresses. **a**, Graphical representations of Col-0 and *p2k1/p2k2* cortex cell PM polarities (digitally magnified from Supplementary Fig. 8a) after fifteen minutes of treatments. **b,** GP values of Col-0 and *p2k1/p2k2* cortex cell PMs. Each data point indicates mean of GP-values per individual root. All statistics were done using one-way ANOVA with Post Hoc – Tukey HSD, *P*-values indicate the statistical significance of differences between treated roots and their respective controls, *alphabet* refers to significance between all samples. Error bar = standard deviation. **c** Digitally magnified image of cortex cells (from Figure 4c) showing (*yellow*) regions of elevated GP values implying elevated Lo lipid presence (*white arrows*) surrounded by Ld lipids. Scale bar = 10 μm **d,** Visualisation showing formation of Lo lipid areas, resulting in decreased overall PM polarity. Formation of these areas increases lipid spacing (indicated by *red dashed* lines and arrow) leads to increased fluidity amongst Ld lipids.

We proceeded to determine frequency shifts and linewidth changes in the cytoplasm and at the cell borders of Col-0 and *p2k1 p2k2* cortex cells mock or 0.3M sorbitol treated for 15 minutes (Figure 5b). Frequency shifts and linewidth were similar in mock conditions in the cytoplasm and at the cell borders in mock-treated Col-0 and *p2k1 p2k2* cortex cells. Frequency shifts changed in response to sorbitol similarly in all genotypes and areas examined. Linewidth did not change in the cytoplasm compared to mock-treated cells in response to sorbitol. However, there was a slight decrease in linewidth in Col-0 cortex cell borders in response to sorbitol, which seems to be less pronounced in *p2k1 p2k2*.

Next, we investigated how loss of P2K1 and P2K2 affects lateral PM organization in cortex cells exposed to different treatments for 15 minutes and four hours using C-Laurdan (Fig. 6a, b and Supplementary Figs. 8 and 9). *p2k1 p2k2* PMs exhibited elevated GP-values in mock conditions (both with/without DMSO) after 15 minutes, indicating that double mutant PMs are less polar (Fig. 6a, b). Col-0 and *p2k1 p2k2* PMs became less polar in response to sorbitol, indicating that P2K1 P2K2 are not required for sorbitol-induced increase in PM order. In response to ISX treatments Col-0 PMs also became less polar but *p2k1 p2k2* GP values did not change. This suggests that P2K1 P2K2 are required to translate an ISX-induced effect on the cell wall to the PM. Lateral organization of Col-0 and *p2k1 p2k2* PMs was similar after four hours under mock conditions (Supplementary Fig. 9). If DMSO is included in the mock treatment, *p2k1 p2k2* PMs were more polar. Col-0 and *p2k1 p2k2* PMs were less polar after sorbitol, ISX, and combined treatments. These results indicate that P2K1 P2K2 activity is required for lateral organization of the PM under mock (DMSO) conditions and the short-term (15 minutes) response to ISX- induced CWD but not long-term (four hours) adaptation to both stresses.

## Discussion

We used molecular rotors (BODIPY type), C-Laurdan and Brillouin microscopy to investigate how the mechanical properties of PM, cell wall, cytoplasm and vacuole change in Arabidopsis roots exposed to hyperosmotic as well as cell wall stress and to establish the role of P2K1 P2K2 kinases in this context. Amongst our findings, three stand out. The PM and cell borders respond within minutes to stress while wall porosity seems to change comparatively little and the vacuole exhibits changes only after several hours. Both ISX and sorbitol affect PM lipid order (based on C-Laurdan results) and fluidity (based on N+-BDP results) with the effects being stress-specific and exhibiting different temporal dynamics. P2K1 P2K2 mediate PM mechanostasis as well as the hyperosmotic stress effect on PM fluidity and the ISX effect on lipid order.

Using N+-BDP we found that both hyperosmotic stress and ISX affect PM fluidity across all tissue layers examined with the extent of the effects influenced predominantly by developmental stage (Figs. 1 and 3). Previously cell layer specific responses to hyperosmotic stress and CWD have been reported ^53–56^. Our results suggest that developmental stages have more pronounced effects on PM fluidity during exposure to hyperosmotic stress or CWD than position in the tissue (outer vs. inner cell layer). Recently we showed using Brillouin microscopy and cell wall-specific molecular rotors that mechanical properties do differ *in vivo* between developmental stages in the cell layers of Arabidopsis root tips ^52^. These observations suggest that PM fluidity changes in response to hyperosmotic stress ^57^ and CWD in a developmental-stage-dependent manner and is possibly influenced by the mechanical properties of the cell walls of the different stages ^52^.

The Brillouin microscopy-based analysis detected hyperosmotic stress-induced changes in frequency shift at the cell borders and in the cytoplasm after fifteen minutes, while linewidth did not change significantly (Fig. 2d). This implies fast responses to both treatments, which started before. Mock treatments affected frequency shift in the opposite direction compared to hyperosmotic stress, suggesting that fresh media affects the mechanical properties in a different way than hyperosmotic stress. These temporal dynamics are aligned with previous research describing how fast turgor changes in response to hyperosmotic stress ^57^. One possible explanation for the results is that the sudden drop in turgor changes water content, solute composition, or apoplastic fluid at the borders leading to a frequency shift change. Alternatively, it could involve controlled modifications of the cell wall structure through activation of CWI signalling and cell wall modifying proteins such as expansins ^58^.

We used N+-BDP to investigate PM fluidity and C-Laurdan to characterize PM order in cortex cells. Decreased lifetimes for N+-BDP indicate increased fluidity while higher GP values for C-Laurdan indicate a less polar, more ordered PM. Hyperosmotic stress raised GP values at early timepoints while N+-BDP lifetimes were reduced, indicating that the PM became more ordered and fluid. In both cases the effects became less pronounced over time. Previous work indicates that hyperosmotic stress induces nanodomain formation in plants ^7,59^ while decreased PM tension can lead to lipid phase separation ^4^. The opposite changes in GP and lifetime suggest that hyperosmotic stress results in ordered nanodomain formation (increased order detected by C-Laurdan) causing the PM matrix surrounding the nanodomains being more loosely organized (more fluid Fig. 6c, d). Alternatively, the increase in fluidity may be caused by PM tension being elevated at PM-cell wall adhesion sites ^21^. These adaptive changes could be mediated by RBOH NADPH oxidases, aquaporins (such as PLASMA MEMBRANE INTRINSIC PROTEIN 2;7) and FER ^4,59,60^. FER is a PM-localized, receptor kinase which is connected to the cell wall and modulates organisation of phosphatidylserine, which in turn influences ROP6-containing nanodomains ^61^. The relevance of FER is further supported by RALF peptides having been implicated in short term induction of lipid-ordered phases ^62^. Other relevant molecular components could be the HYPERSENSITIVE-INDUCED REACTION2 (HIR2) stomatin/prohibitin/ flotillin/HflK/ C (SPFH) family protein or REMORINs. REMORINs bind anionic phospholipids, mediate PM nano-organisation and have been implicated in PM-cell wall attachment during exposure to hyperosmotic stress ^20,63,64^. HIR2 has been implicated in formation of PM sub-compartments and could provide a structural framework for spatial PM organization^65^.

We also analysed the impact of ISX and ISX/sorbitol cotreatments on the mechanical properties of the PM (Fig. 4). Since the routinely used ISX solvent DMSO affects mechanical properties of the PM (GP values changed after 15 minutes) we are interpreting ISX and ISX/sorbitol results relative to DMSO. ISX-treated PMs exhibited pronounced GP increases before any effects were observed with N+-BDP or Brillouin microscopy (Fig. 4a, d, e). A potential cause for this rapid effect on PM order could be PM detachment from the cell wall due to loss of cellulose synthase complexes linking PM and cell wall. Such detachment can be caused by stress ^20^ or ISX ^66^. Lifetime of the cell wall localised CarboTag-BDP dipped between one to three hours while N+-BDP lifetime fell after three hours to a level observed in sorbitol-treated PMs. A possible explanation is that the altered cell wall yields to turgor pressure, consequently the PM is exposed to enhanced load and tension increases. In PMs exposed to combined ISX/sorbitol treatments the early transient lifetime dip of N+-BDP, observed in response to sorbitol, was absent. Since we did not detect an effect on the cell border (Brillouin microscopy) or cell wall porosity (CarboTag-BDP) a plausible explanation is that this effect is caused by previously described ISX-induced alterations of the PM-cell wall linkage ^20^.

Interestingly the ISX-induced CarboTag-BDP lifetime changes between one and three hours are reduced in ISX/sorbitol-treated cells, possibly due to a reduction in turgor. This aligns well with previously reported Osmo sensitivity of ISX-induced responses ^3,51,55^.

The PM-specific molecular rotor shows that the PM becomes more fluid at the time points where the cell wall-specific rotor detected only minor CW porosity changes. Intriguingly, combined ISX/sorbitol treatments suppress both early PM fluidity changes induced by sorbitol alone as well as later changes in cell wall porosity caused by ISX alone. Results from the C-Laurdan-based analysis of PM organization indicate decreased PM polarity caused by individual treatments (DMSO, ISX, and sorbitol), while the effect is less pronounced in PMs exposed to combined sorbitol/ISX treatments. These observations suggest that the PM-cell wall continuum is reorganized in response to qualitatively different forces and/or stimuli generated by ISX/CWD and hyperosmotic stress (Fig. 6b, c). These changes may affect PM-cell wall connections, involve formation of novel/reorganisation of existing nanodomains or modify PM lipid composition i.e., affect abundance of major sphingolipid types such as GlcCers and GIPCs ^14,16,59,67^. The apparent suppression of the early transient changes in mechanical properties and PM polarity induced by sorbitol/ ISX co-treatments could be explained in two ways. The first explanation would involve only passive adaptation of the PM and cell wall to changes in turgor. The second would involve at least two different, interconnected processes. One mediates adaptation to hyperosmotic stress, and another one mediates adaptation to CWD which in turn can also modulate the hyperosmotic stress-induced process.

Importantly, this is not a one-way road between both processes since previous work has shown that ISX-induced responses such as jasmonic acid induction are suppressed by sorbitol co-treatments ^3,51^.

In plants, different mechanisms have been proposed for detecting disturbance in the PM-cell wall continuum. One mechanism would involve PM tension-sensitive ion channels exemplified by hyperOSmolarity-gated CAlcium-permeable channels (OSCA), which upon sensing PM tension changes, facilitate Ca^2+^-influx into the cytoplasm and initiate adaptive processes including Ca^2+^-dependent PM repair ^10,68^. An alternative mechanism could consist of FER-containing signalling complexes located in nanodomains, which involve reactive oxygen species and/or Ca^2+^-based signalling processes and receptor kinases like FER ^6,7,59^. Additional alternatives implicate cellulose microfibrils in combination with cellulose synthase complexes ^20^ or LecRK-I.9/P2K1 in combination with Hechtian strands ^24^. Manipulating the binding of LecRK-I.9/P2K1 to the cell wall causes changes in the mechanical properties of the wall and the response to hyperosmotic stress, while LecRK-I.9/P2K1 overexpression induces increased formation of Hechtian strands.

Here we find that cell wall porosity and PM fluidity/polarity in *p2k1 p2k2* cortex cells differ from Col-0 in mock conditions. Sorbitol-induced PM fluidity changes were absent in *p2k1 p2k2* cortex cells while GP rises were enhanced early on (Figs. 5a, 6b). Sorbitol-induced changes in cell wall porosity (CarboTag-BDP) and in border frequency shift were similar in both genotypes, whereas the sorbitol-induced reduction in border linewidth was weaker in *p2k1 p2k2* (Fig. 5a, b). The sorbitol-induced change in wall porosity in Col-0 was absent in *p2k1 p2k2* after four hours (Supplementary Fig. 7a). These results suggest that P2K1 P2K2 are required for the mechanostasis of the PM under steady state conditions and modulate the extent of change in PM lateral organization in response to perturbations. Since these perturbations can be brought about by sorbitol, DMSO and ISX P2K1 P2K2 may not modulate responses to a particular stress but are generally required for PM mechanostasis. The mechanical properties of the wall and vacuole are adapted subsequently. Two models could explain the activities of P2K1 P2K2. In the first model P2K1 P2K2 would function as lectin-based PM-cell wall anchors coordinating PM-cell wall interactions but information the mode of action is limited ^24–26^. The second model would involve early release of extracellular ATP in response to sorbitol or CWD ^69^, meaning P2K1 P2K2 act as eATP receptors. They in turn activate signaling processes involving calcium and nanodomain localized ROP6/RBOH ^7,29–31,59^. This model suggests that P2K1 P2K2 activity eventually influences vacuole viscosity, likely as a downstream consequence of its earlier effects at the PM and cell wall rather than directly acting on the vacuole.

To summarize, our results suggest a temporal hierarchy how the micromechanical properties of subcellular structures adapt to hyperosmotic stress and CWD: PM fluidity and polarity respond within minutes, cell wall porosity changes are comparatively modest and structure-dependent, and vacuole viscosity is the last property to change, consistent with an indirect, downstream adaptation rather than a primary stress response. P2K1 P2K2 appear to modulate this hierarchy at multiple points, contributing to early PM adaptation as well as to longer-term cell wall and vacuole responses, without being universally required across all structures and timepoints examined. The sequential, structure-specific timing observed here suggests that sub-cellular structures respond to hyperosmotic stress and CWD on different timescales. It should be noted that these conclusions are based on the *p2k1 p2k2* double mutant. Since single mutants were not examined here, the individual contributions of P2K1 and P2K2 to the described phenotypes remain to be determined. Important questions to address in the future are if/how eATP-based signalling processes involving P2K1 P2K2 and absence/presence of Hechtian strands contribute to the adaptive changes in micromechanical properties described here.

## Methods

### Plant material

*Arabidopsis thaliana* Col-0 accession seeds were obtained from the European Arabidopsis Stock Centre in Nottingham. The *p2k1 p2k2* double mutant is described in ^29^.

### Plant growth

Seeds were sterilized using 70% ethanol, followed by 50% (v/v) bleach (Klorin) for 5 minutes. Then the bleach was discarded and the seeds were washed three times with sterilized water. Sterile seeds were sown on solid media plate pH 5.7 containing ½ Murashige and Skoog (Merck, M5519-50L), 0.05% MES sodium salt (Merck, M3058-1KG), 1% sucrose (Merck, S7903-5KG), and 1% agar (Merck, A1296-1KG). The plates were stratified for 24-48 hours at 4°C before transferring them into the growth cabinet. Seeds were then grown under long-day conditions (16-hour light/8- hour dark, 150 μmol⋅m^−2^⋅s^−1^ photon flux density, 21°C, and 40% humidity) for 6 days (counting the next day as the first day). Seeds germinated on the second day after transfer, hence all seedlings used in this study are 4-DAG (4 days after germination) unless stated otherwise.

### Plant treatment

Seedlings with approximately the same root length were selected and transferred into in cell culture disks (VWR, 734-2814) containing liquid media with either 0.3M sorbitol (d-sorbitol; Merck, 56021-5KG), 0.6μM ISX (Merck, 75772-50MG; 600μM stock was dissolved in DMSO), a combined treatment of sorbitol and ISX, 14.1mM DMSO (Merck, D8418-50ML), and a liquid media without any treatment as a control. We used a ½ MS liquid media as a control for the sorbitol treatment, while DMSO served as a control/mock-treatment for the ISX and sorbitol + ISX treatments. The liquid media used for treatments, staining, and mounting were prepared according to the solid media, excluding agar and sucrose. Seedlings were then treated for 5 mins, 15 mins, 30 mins, 1h, 2h, 3h, and 4h. The treatment durations were included in the total treatment times. For example, to study roots after 4 hours of hyperosmotic stress with a dye that requires 1 hour of staining, seedlings were first treated with sorbitol-containing medium for 3 hours. Subsequently, the treatment solution was replaced with sorbitol-containing medium with the dye for 1 hour. For shorter treatment durations, such as 15 minutes, seedlings were first treated with the staining solution in a liquid medium for 45 minutes. Then transferred to medium containing both dye and sorbitol for the last 15 minutes. This protocol was consistently applied to all treatments and staining procedures in the study.

### Fluorescence lifetime imaging (FLIM) for microviscosity

Seedlings were stained with 20μM CarboTag-BDP (cell wall), 20μM N+-BDP (plasma membrane), and 10μM Sulfo-BDP (vacuole) in a liquid medium without sucrose for 2 hours, 30 minutes, and 3 hours ^34,35^. Seedling were briefly washed to remove the excess probe and then mounted in a liquid medium. FLIM was performed using a Leica SP8 SMD/MD (Leica Microsystems) with an HCX IRAPO L 25x/0.95 water-immersion objective, and 512 × 512 pixels images were taken with a line scanning speed of 600Hz in unidirectional mode, a pinhole of 1 airy unit, and 1× digital zoom. The signal was detected using a single-photon avalanche photodiode (APD) following the configuration: CarboTag-BDP (white light laser [WLL] 488nm, detector gain: 200%, laser: 40%); N^+^-BDP (WLL 514nm, detector gain: 100%, laser: 5-8%); Sulfo-BDP (WLL 514nm, detector gain: 250%, laser: 20%). The photon was then collected using a time-correlated single-photon counting (TCSPC) module PicoHarp 300 (Picoquant gmbh, Germany) with image acquisition period of 100.15 seconds at a repetition rate of 20 MHz.

FLIM analysis was performed using SymphoTime 64 (PicoQuant, Germany). To measure plasma membrane fluidity across different tissues and root zones (Fig. 1 and 3), images were taken from the focal plane in the center of the root where all tissues were visible, and ROIs for each tissue were manually selected (Supplementary Fig. 2a). For cortex-specific measurements, we adjusted the microscopy depth so that majority cortex cells were visible, then manually selected cortex ROIs, excluding areas with ectopic probe internalization (Supplementary Fig. 2b). Then the fitting was done using n-exponential tailfit model with the model parameter set to *n* = 2. The area of measurement (*x* axis) in the TCSPC graph was adjusted to start from 0.5 ns after the instrument response function (IRF) until 1 ns before the end-tail of the decay curve and the histogram was recalculated. The histogram was then used to adjust the false colour scale to generate the fluorescence lifetime maps which represent the mean fluorescence lifetime per pixel in nanoseconds. In terms of the fluorescence lifetime maps of roots (Fig. 2A), the images were further stitched to generate a composite image of the whole roots using the MosaicJ plugin in Fiji ^70^. Furthermore, the intensity-weighted average lifetime (Tau_Av_Int) values were plotted over time. Each data point represents the mean lifetime of multiple cells from a single tissue per root, with measurements conducted on 8 to 10 roots for each timepoint and treatment.

### Lipid polarity imaging

Seedlings were stained with 40μM C-Laurdan (BioTechne, 7273) for 1 hour. After staining, seedlings were briefly washed before being mounted in liquid medium. Fluorescence images of 1024 x 1024 pixels were obtained using an inverted Leica SP8 SMD/MD with an HCX IRAPO L 25x/0.95 water-immersion objective, unidirectional scan speed of 400Hz, a pinhole of 1 airy unit, and 1× digital zoom. The probe was excited using two-photon laser at 780 nm, with 10% intensity, 60% gain, and 12% offset. Fluorescence was detected using HYD1 detectors with a range of 415 - 455nm and gain: 203.8%, and HYD2 detectors with a range of 490 - 530nm and 200.3%. Image analysis was performed in Fiji, where the ROIs (plasma membrane) were manually selected. Then the mean generalized polarization (GP) values were analysed using fluidity calculator v2.0 macro tool (available at: https://github.com/pcarravilla/fluiditycalculator) ^71^ and plotted in a graphic. Each data point corresponds to the mean of per-pixel GP values, calculated from multiple selected cells within a single tissue per root. Measurements were obtained from 7 to 11 roots for each genotype and treatment. Additionally, the jet-black-minimum LUT was used to visualize the GP image (available at: https://github.com/quokka79/GPcalc).

### Data processing and statistical analysis

Data were plotted in R using the ggplot2 package and the statistical analysis was done using IBM SPSS Statistics ^72^. Statistics were done using either one- or two-way ANOVA with Post Hoc – Tukey HSD corrections or Welch’s t-test with Holm correction. In Fig. 5a and Supplementary Fig. 7, we used for wildtype combined datasets in which each quadratic and rectangular cortical cell was plotted as an individual data point; thus, the graph shows twice the number of data points per biological replicate.

### Brillouin microscopy

The study was conducted using the custom-built Brillouin module, which includes an inverted Leica SP8 confocal microscope coupled to a two-stage VIPA Brillouin spectrometer ^52^. Excitation was provided by a 532 nm laser (Cobolt), which was directed to the backport of the confocal microscope. A 40X NA 1.1 water immersion objective was used to illuminate the sample, and the light was collected in a backscattering configuration. The scattered light was focused to a single-mode optical fibre before entering the spectrometer, which consisted of two-stage VIPA etalons with a 30 GHz free spectral range (Light Machinery). A qCMOS camera was used to collect the spectra.

Image acquisition was set at 100 ms and laser power to 10-15 mW. Point-by-point imaging was performed with a 1 μm distance between adjacent pixels in the X-Y plane and a frame integration of four images. Samples were imaged with transmitted-light wide-field illumination before and after recording the Brillouin spectra. No phototoxicity effects were observed. A custom MATLAB (Mathworks) script was used to extract the Brillouin peak positions and linewidth after fitting with a Lorenztian function^52^. The pixels were ordered according to the stage position to generate heatmaps of frequency shift and linewidth. The values of frequency shift and linewidth are relative to values measured in media. Water and methanol were regularly measured as reference samples to calibrate the frequency axis. An overlap of the transmitted-light and Brillouin images was done manually to define the region of interest (border and area).

## Acknowledgement

We gratefully thank Astrid Bjørkøy from the Center for Advanced Microscopy, Department of Physics, Norwegian University of Science and Technology for her technical assistance. We thank members of the Hamann lab for constructive input on the manuscript and Daniela Sueldo, Leah Gachao for providing *p2k1 p2k2* seeds.

## Author information

Dhika Amanda, Luis Alonso Baez & Thorsten Hamann

1. Department of Biology, Norwegian University of Science and Technology, Hogskoleringen 5, Trondheim 7491, Norway Óscar A. Badias
2. Universitat Politècnica de València (UPV), Calle de Ramiro de Maeztu, 46022, València, Spain Joris Sprakel & Maarten Besten
3. Laboratory of Biochemistry, Wageningen University & Research, Stippeneng 4, 6708 WE, Wageningen, The Netherlands

## Contributions

D.A: conceptualized the project, designed the experiments, performed majority of the experiments, analysed and visualized data, co-supervised O.A.B, acquired funding and project administration, wrote the original draft. O.A.B: conducted vacuole viscosity experiments and analysis. J.S and M.B: provided the resources for molecular rotors and methodology, edited the manuscript. L.A.B: performed Brillouin microscopy experiments and data analysis, visualization, methodology. T.H: conceptualized the project, provided supervision, acquired funding and contributed to project administration, wrote the original draft, and edited the manuscript together with the coauthors.

## Data availability

The manuscript does not contain any data that meets the mandatory deposition requirement as described (https://www.nature.com/ncomms/editorial-policies/reporting-standards). All data produced here will be made available by the corresponding author, Thorsten Hamann, upon reasonable request.

## Ethics declarations

**Supplementary Figure 1. Cell wall porosity, plasma membrane fluidity, and vacuole crowding of cortex quadratic and rectangular cells in the elongation zone of Col-0 roots exposed to hyperosmotic stress**. Mean fluorescence lifetimes of N+-BDP (PM: plasma membrane), CarboTag-BDP (CW: cell wall) and sulfo-BDP (VC: vacuole) in mock- or 0.3M sorbitol-treated Col-0 roots. X axis indicates treatment period. Y axis indicates average fluorescence lifetime in nanoseconds (ns). From left to right, respectively, quadratic cells: n control-PM = 10, 8, 10, 8, 9, 8, 10, 8; all n for treated-PM = 10, except 4h = 9; all n for control-CW = 9, 10, 10, 10, 10, 10, 9, 10; n treated-CW = 9, 10, 10, 10, 10, 10, 10, 8; all n VC = 10, except control 4h = 8 and sorbitol 4h = 11. Rectangular cells: n control-PM = 10, 8, 10, 8, 9, 8, 10, 8; all n treated- PM = 10; all n CW = 10, except 3h control = 9 and 4h treatment = 8; all n VC = 10, except control 4h = 8 and sorbitol 4h = 12. Error bars = standard deviation. Statistics: one-way ANOVA with Post Hoc – Tukey HSD (*asterisk* refers to significance between treated roots vs control at each timepoint, *alphabet* refers to significance between timepoints within one sample groups, ¤ refers to significance between the cell shape, ¤ = control group and ¤ = treated group).

**Supplementary Figure 2. Fluorescence lifetime maps of the cell wall (CarboTag-BDP), plasma membrane (N+-BDP), and vacuole (sulfo-BDP) in Col-0 root elongation zones.** a, Example of ROI selections of cells in different tissues. b, Example of ROI selections of cortex cells in samples without- and with probe internalization. c, Representative FLIM images of Col-0 roots after mock or 0.3M sorbitol treatments. Scale bar = 100μm. *Arrows* indicate ectopic probe accumulation.

**Supplementary Figure 3. Fluorescence lifetime maps of the cell wall (CarboTag-BDP), plasma membrane (N+-BDP), and vacuole (sulfo-BDP) in Col-0 root elongation zones in response to CWD**. Representative FLIM images of Col-0 roots after mock (DMSO) or 0.6μM ISX treatments. Scale bar = 100μm. *Arrows* indicate ectopic probe accumulation.

**Supplementary Figure 4. FLIM maps of the cell wall (CarboTag-BDP), plasma membrane (N+-BDP), and vacuole (sulfo-BDP) in Col-0 root elongation zones exposed to hyperosmotic stress and CWD**. Representative FLIM images of Col-0 roots after mock (DMSO) or 0.3M sorbitol/0.6μM ISX treatments. Scale bar = 100μm. *Arrows* indicate ectopic probe accumulation.

**Supplementary Figure 5. Fluorescence lifetime maps of the cell wall (CarboTag-BDP), plasma membrane (N+-BDP), and vacuole (sulfo-BDP) in *p2k1/p2k2* root elongation zones after 15 minutes of hyperosmotic stress or CWD**. Representative FLIM images of *p2k1/p2k2* roots after control, mock (DMSO), 0.3M sorbitol, 0.6μM ISX treatments. Scale bar = 100μm.

**Supplementary Figure 6. Fluorescence lifetime maps of the cell wall (CarboTag-BDP), plasma membrane (N+-BDP), and vacuole (sulfo-BDP) in root elongation zones after 4 hours long treatments**. Representative FLIM images of *p2k1/p2k2* roots after control, mock (DMSO), 0.3M sorbitol, 0.6μM ISX or combined treatments. Scale bar = 100μm.

**Supplementary Figure 7. Cell wall porosity, plasma membrane fluidity, and vacuole crowding of wildtype and *p2k1/p2k2* cortex cells after four hours. Mean fluorescence lifetime of cell wall (a)**, plasma membrane **(b)**, and vacuole **(c)** in seedlings treated with of control, mock (DMSO), 0.3M sorbitol, 0.6μM ISX, and combination of both. Error bars = standard deviation. Statistics: one-way ANOVA with Post Hoc – Tukey HSD (*P*-values indicate the statistical significance of differences between treated roots and their respective control or DMSO groups, *alphabet* refers to significance between all samples).

**Supplementary Figure 8. Plasma membrane polarity pattern of root elongation zones in Col-0 and *p2k1/p2k2* seedling roots after (a) fifteen minutes or (b) four hours treatments.** Representative images of roots stained with C-Laurdan after control, mock (DMSO), 0.3M sorbitol/0.6 μM ISX or combined treatments. Scale bar = 100μm.

**Supplementary Figure 9. Plasma membrane polarity pattern in cortex cells of Col-0 and *p2k1/p2k2* root elongation zones**. **a,** Graphical representations of wildtype and *p2k1/p2k2* cortex cells (digitally magnified from Supplementary Fig. 8b) stained with 40μM C-Laurdan after control, mock (DMSO), 0.3M sorbitol, 0.6μM ISX, and combined treatments for four hours. **b,** GP values of plasma membrane of wild-type and *p2k1/p2k2* cortex cells. Each data point indicates mean of cortex cell GP-values per individual root. Error bars = standard deviation. Statistics: one-way ANOVA with Post Hoc – Tukey HSD (*P*-values indicate the statistical significance of differences between treated roots and their respective control or DMSO group, *alphabet* refers to significance between all samples).

