## Supplemental FIgure 1-9 for "P2K1 P2K2 are required for early, mechanical adaptation to osmotic and cell wall stress in Arabidopsis roots"

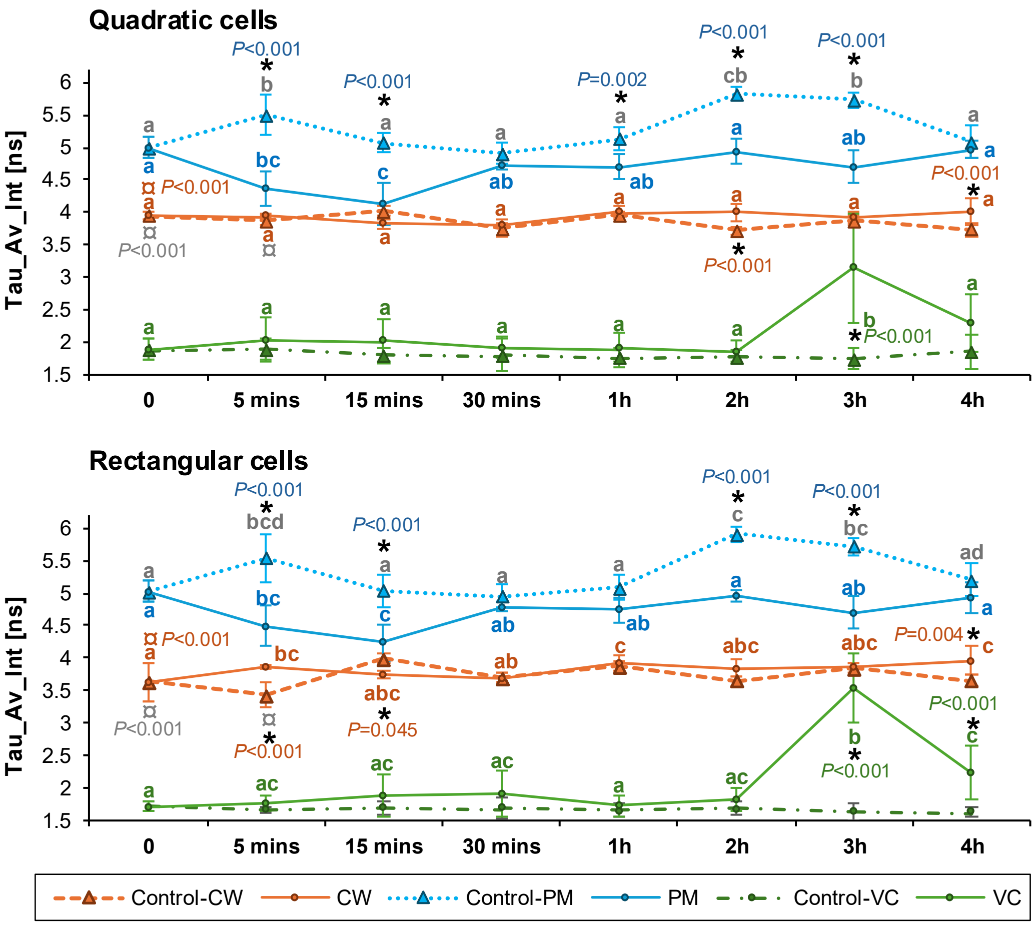


**Supplementary Figure 1. Cell wall porosity, plasma membrane fluidity, and vacuole crowding of cortex quadratic and rectangular cells in the elongation zone of Col-0 roots exposed to hyperosmotic stress**. Mean fluorescence lifetimes of N+-BDP (PM: plasma membrane), CarboTag-BDP (CW: cell wall) and sulfo-BDP (VC: vacuole) in mock- or 0.3M sorbitol-treated Col-0 roots. X axis indicates treatment period. Y axis indicates average fluorescence lifetime in nanoseconds (ns). From left to right, respectively, quadratic cells: n control-PM = 10, 8, 10, 8, 9, 8, 10, 8; all n for treated-PM = 10, except 4h = 9; all n for control-CW = 9, 10, 10, 10, 10, 10, 9, 10; n treated-CW = 9, 10, 10, 10, 10, 10, 10, 8; all n VC = 10, except control 4h = 8 and sorbitol 4h = 11. Rectangular cells: n control-PM = 10, 8, 10, 8, 9, 8, 10, 8; all n treated-PM = 10; all n CW = 10, except 3h control = 9 and 4h treatment = 8; all n VC = 10, except control 4h = 8 and sorbitol 4h = 12. Error bars = standard deviation. Statistics: one-way ANOVA with Post Hoc – Tukey HSD (*asterisk* refers to significance between treated roots vs control at each timepoint, *alphabet* refers to significance between timepoints within one sample groups, ¤ refers to significance between the cell shape, ¤ = control group and ¤ = treated group).

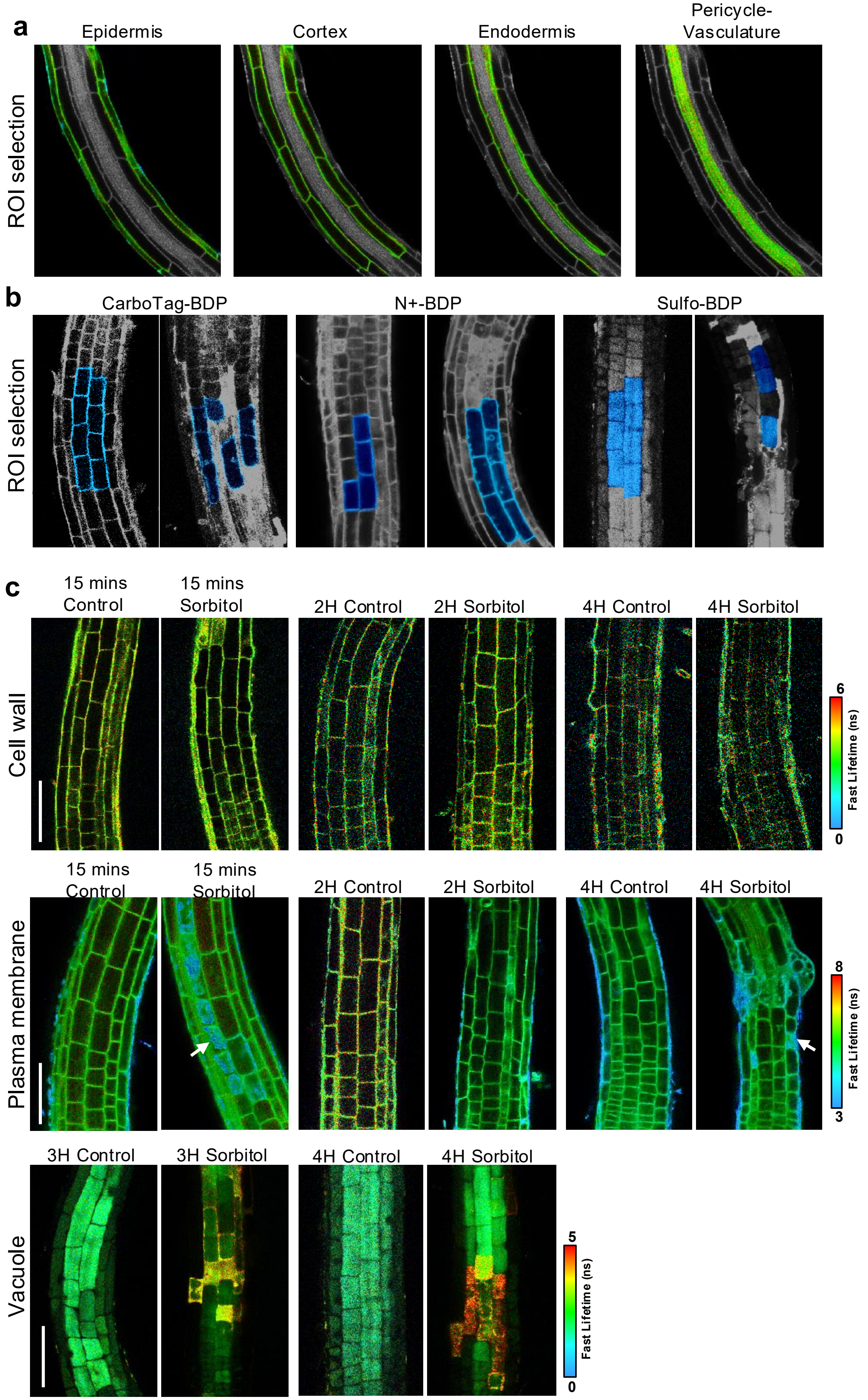


**Supplementary Figure 2. Fluorescence lifetime maps of the cell wall (CarboTag-BDP), plasma membrane (N+-BDP), and vacuole (sulfo-BDP) in Col-0 root elongation zones. a,** Example of ROI selections of cells in different tissues. **b,** Example of ROI selections of cortex cells in samples without- and with probe internalization. **c,** Representative FLIM images of Col-0 roots after mock or 0.3M sorbitol treatments. Scale bar = 100µm. *Arrows* indicate ectopic probe accumulation.

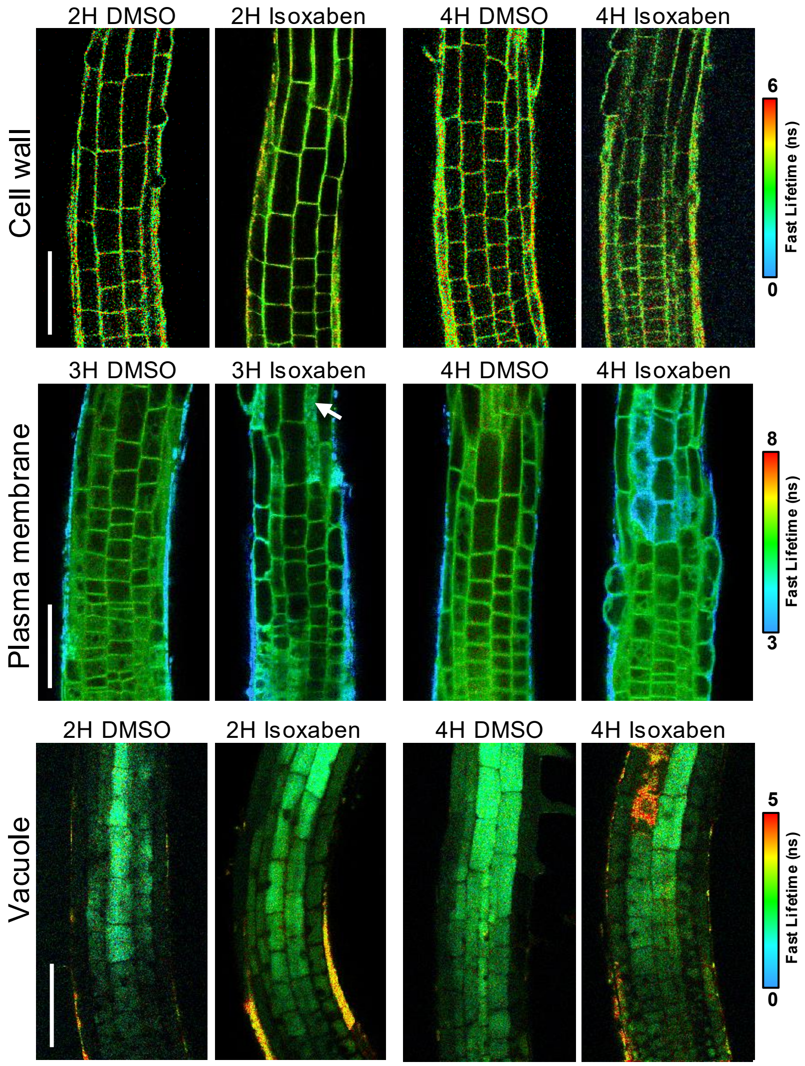


**Supplementary Figure 3. Fluorescence lifetime maps of the cell wall (CarboTag-BDP), plasma membrane (N+-BDP), and vacuole (sulfo-BDP) in Col-0 root elongation zones in response to CWD**. Representative FLIM images of Col-0 roots after mock (DMSO) or 0.6μM ISX treatments. Scale bar = 100μm. *Arrows* indicate ectopic probe accumulation.

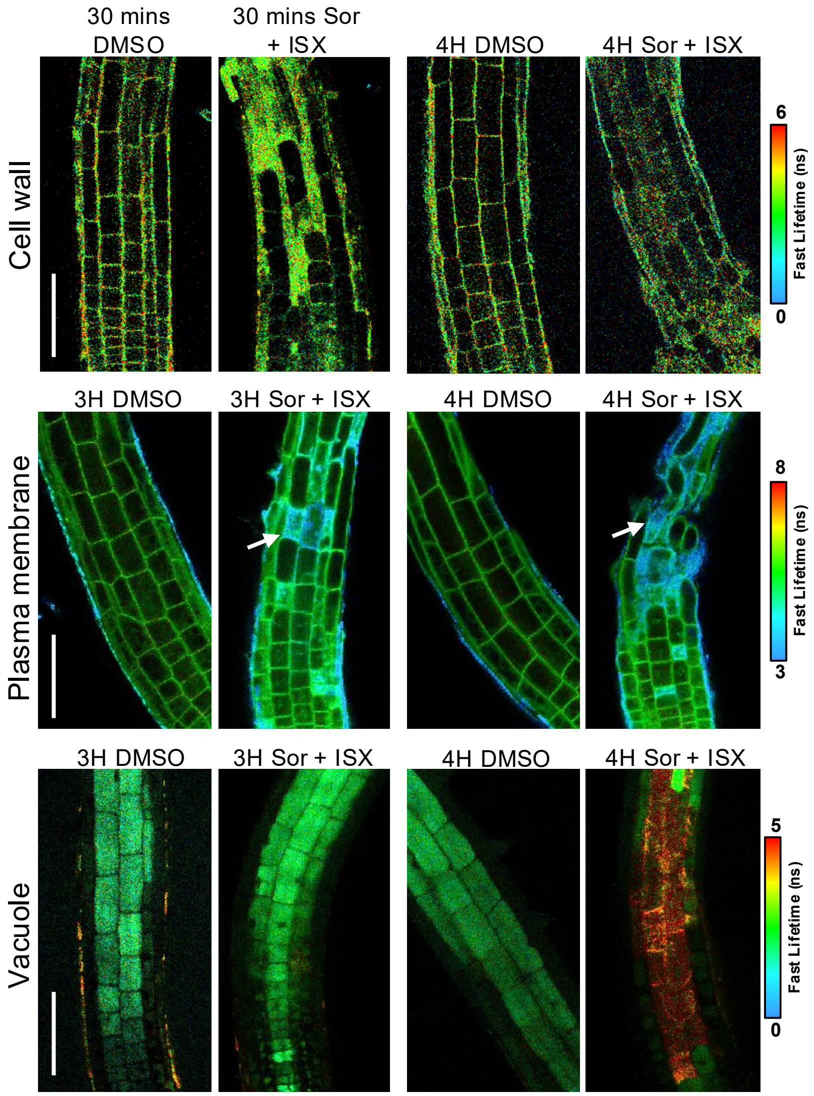


**Supplementary Figure 4. FLIM maps of the cell wall (CarboTag-BDP), plasma membrane (N+-BDP), and vacuole (sulfo-BDP) in Col-0 root elongation zones exposed to hyperosmotic stress and CWD**. Representative FLIM images of Col-0 roots after mock (DMSO) or 0.3M sorbitol/0.6μM ISX treatments. Scale bar = 100μm. *Arrows* indicate ectopic probe accumulation.

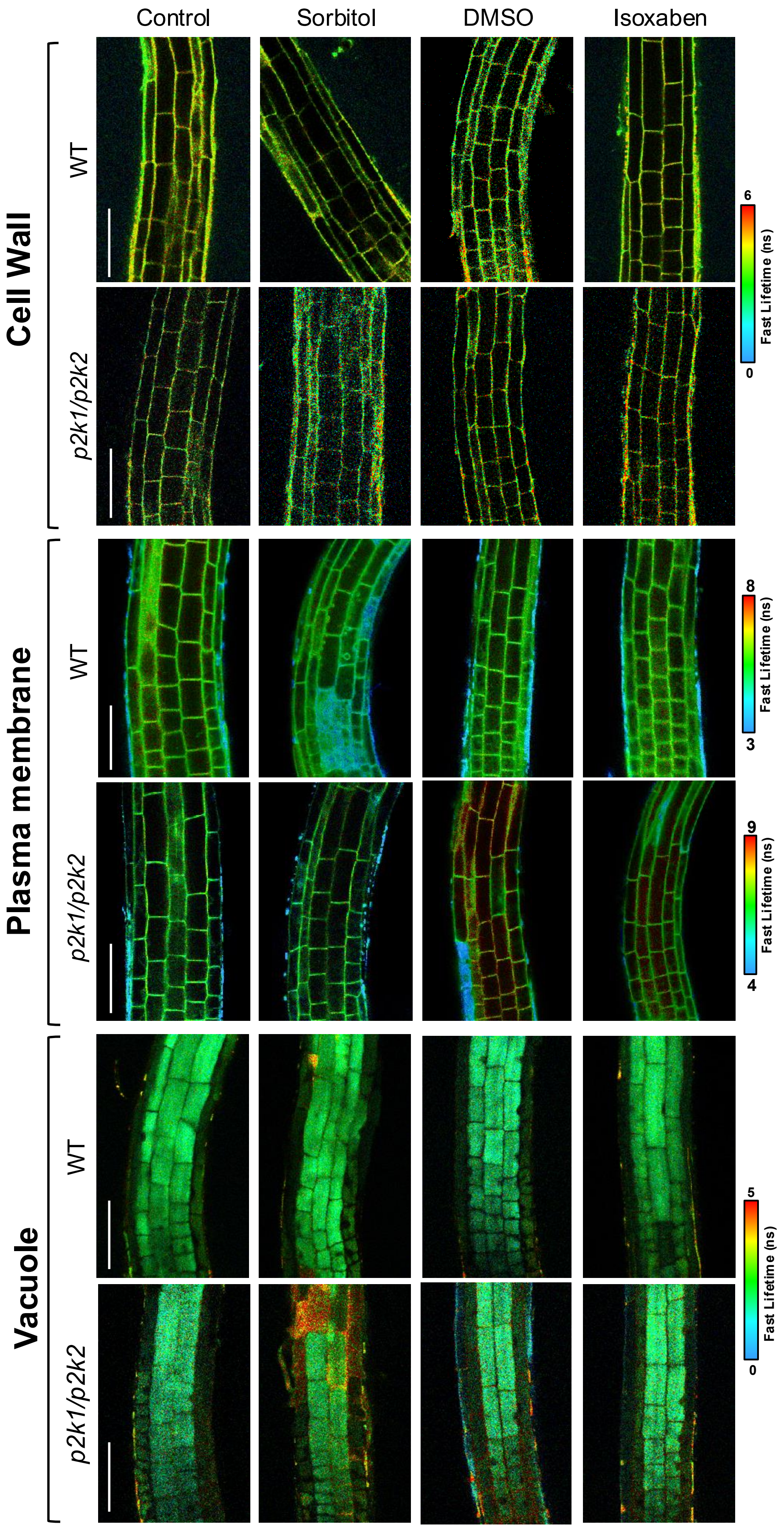


**Supplementary Figure 5. Fluorescence lifetime maps of the cell wall (CarboTag-BDP), plasma membrane (N+-BDP), and vacuole (sulfo-BDP) in *p2k1/p2k2* root elongation zones after 15 minutes of hyperosmotic stress or CWD**. Representative FLIM images of *p2k1/p2k2* roots after control, mock (DMSO), 0.3M sorbitol, 0.6μM ISX treatments. Scale bar = 100μm.

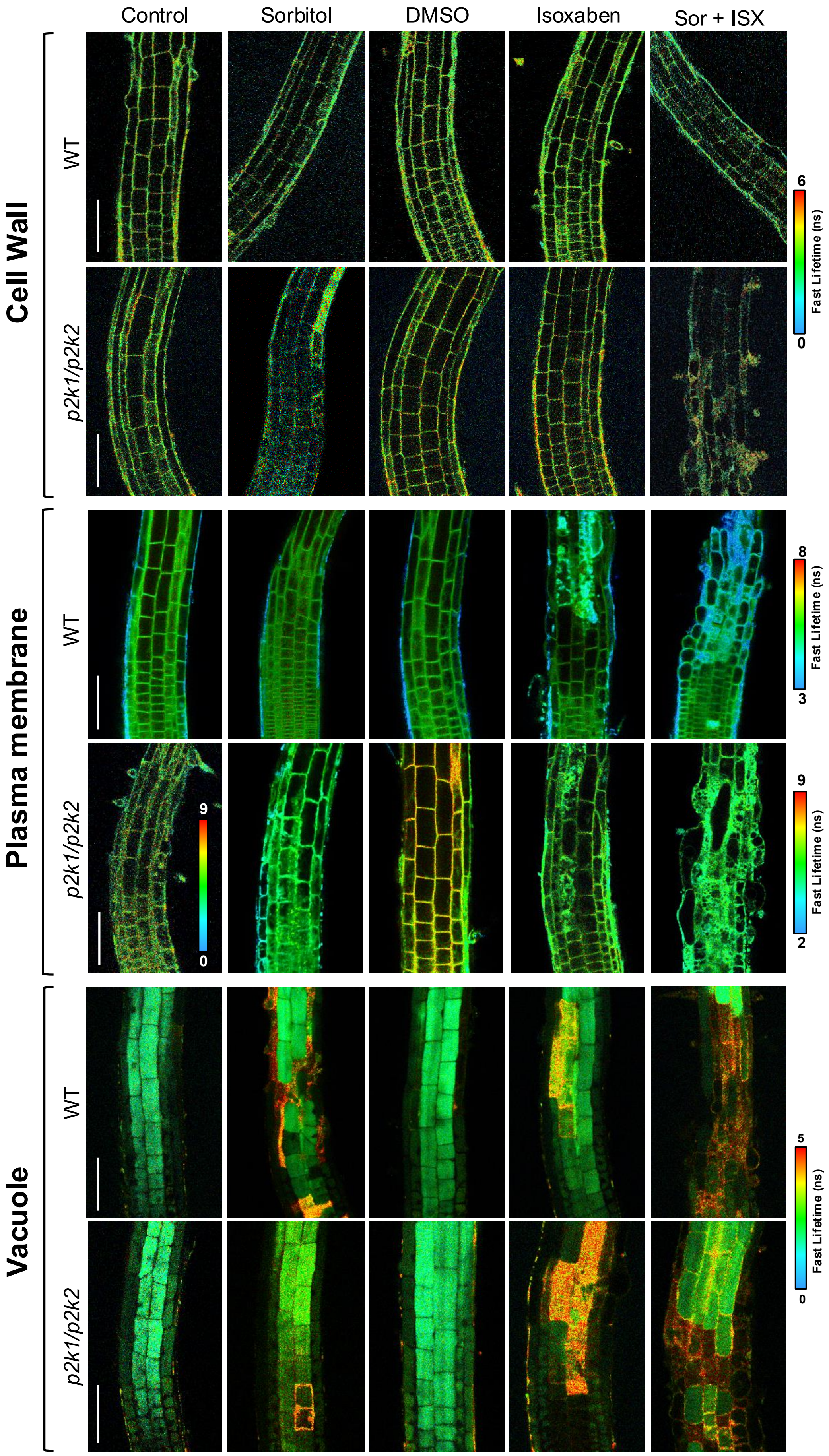


**Supplementary Figure 6. Fluorescence lifetime maps of the cell wall (CarboTag-BDP), plasma membrane (N+-BDP), and vacuole (sulfo-BDP) in root elongation zones after 4 hours long treatments**. Representative FLIM images of *p2k1/p2k2* roots after control, mock (DMSO), 0.3M sorbitol, 0.6μM ISX or combined treatments. Scale bar = 100μm.

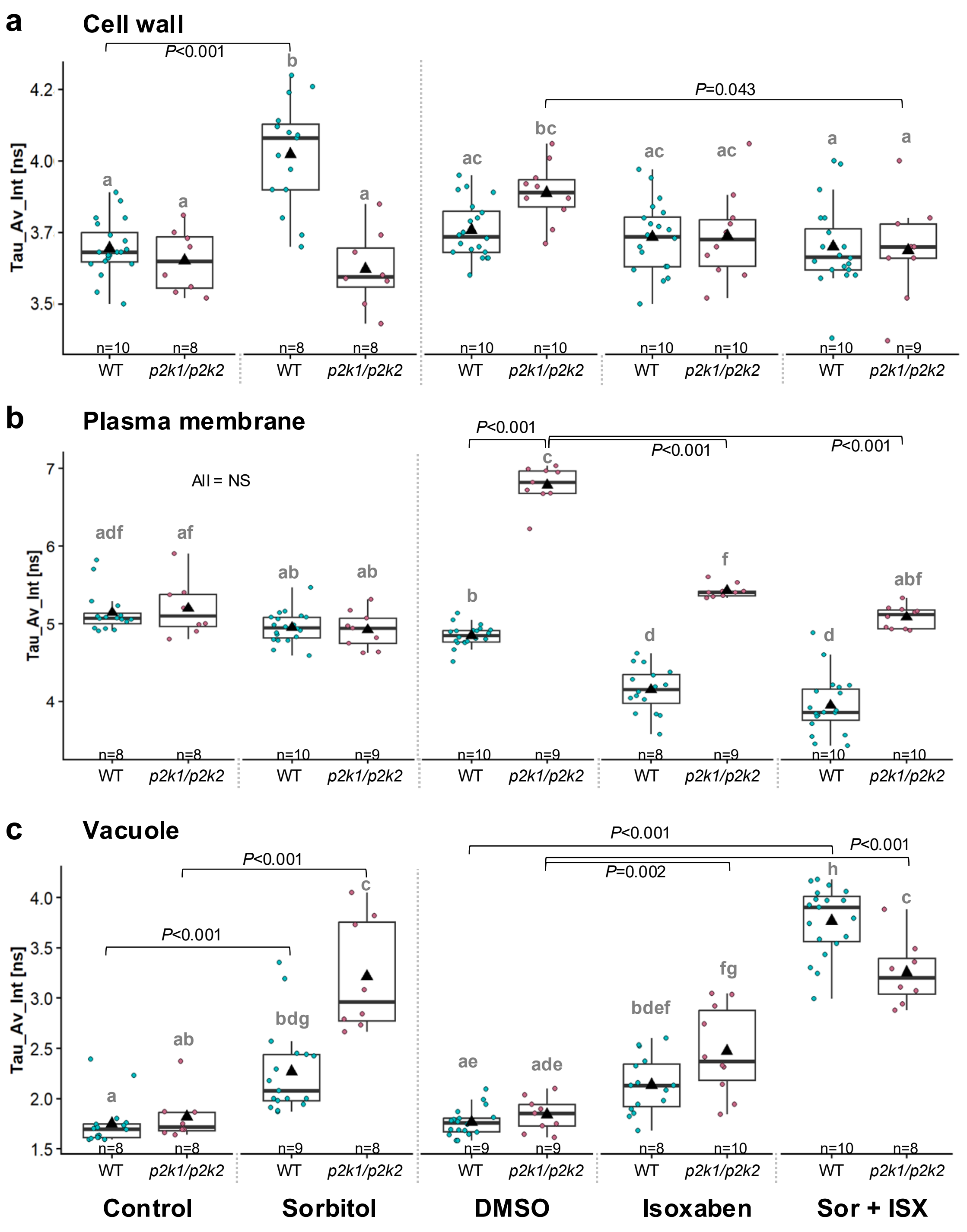


**Supplementary Figure 7. Cell wall porosity, plasma membrane fluidity, and vacuole crowding of wildtype and *p2k1/p2k2* cortex cells after four hours.** Mean fluorescence lifetime of cell wall **(a)**, plasma membrane **(b)**, and vacuole **(c)** in seedlings treated with of control, mock (DMSO), 0.3M sorbitol, 0.6µM ISX, and combination of both. Error bars = standard deviation. Statistics: one-way ANOVA with Post Hoc – Tukey HSD (*P*-values indicate the statistical significance of differences between treated roots and their respective control or DMSO groups, *alphabet* refers to significance between all samples).

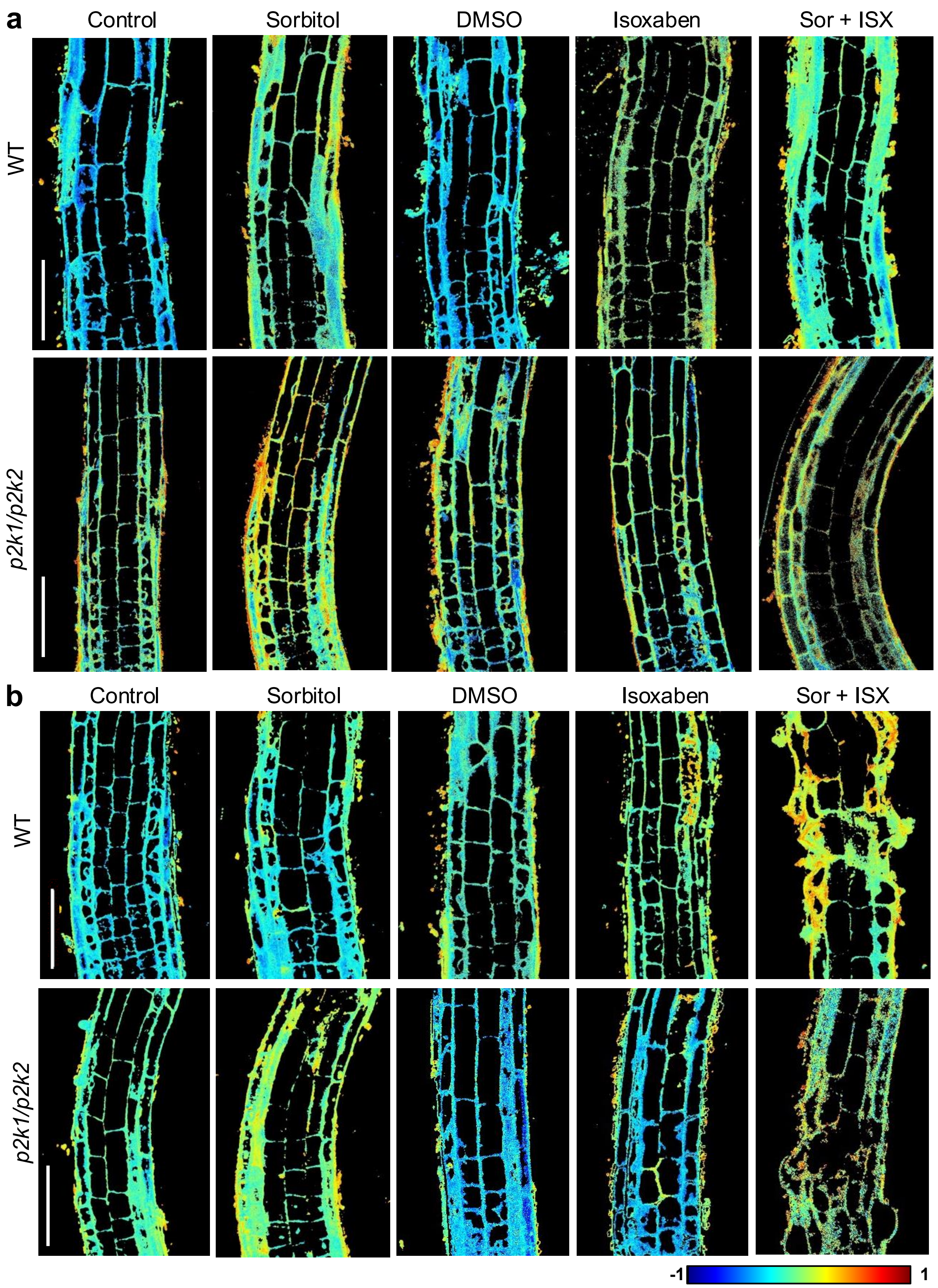


**Supplementary Figure 8. Plasma membrane polarity pattern of root elongation zones in Col-0 and *p2k1/p2k2* seedling roots after (a) fifteen minutes or (b) four hours treatments**. Representative images of roots stained with C-Laurdan after control, mock (DMSO), 0.3M sorbitol/0.6 μM ISX or combined treatments. Scale bar = 100μm.

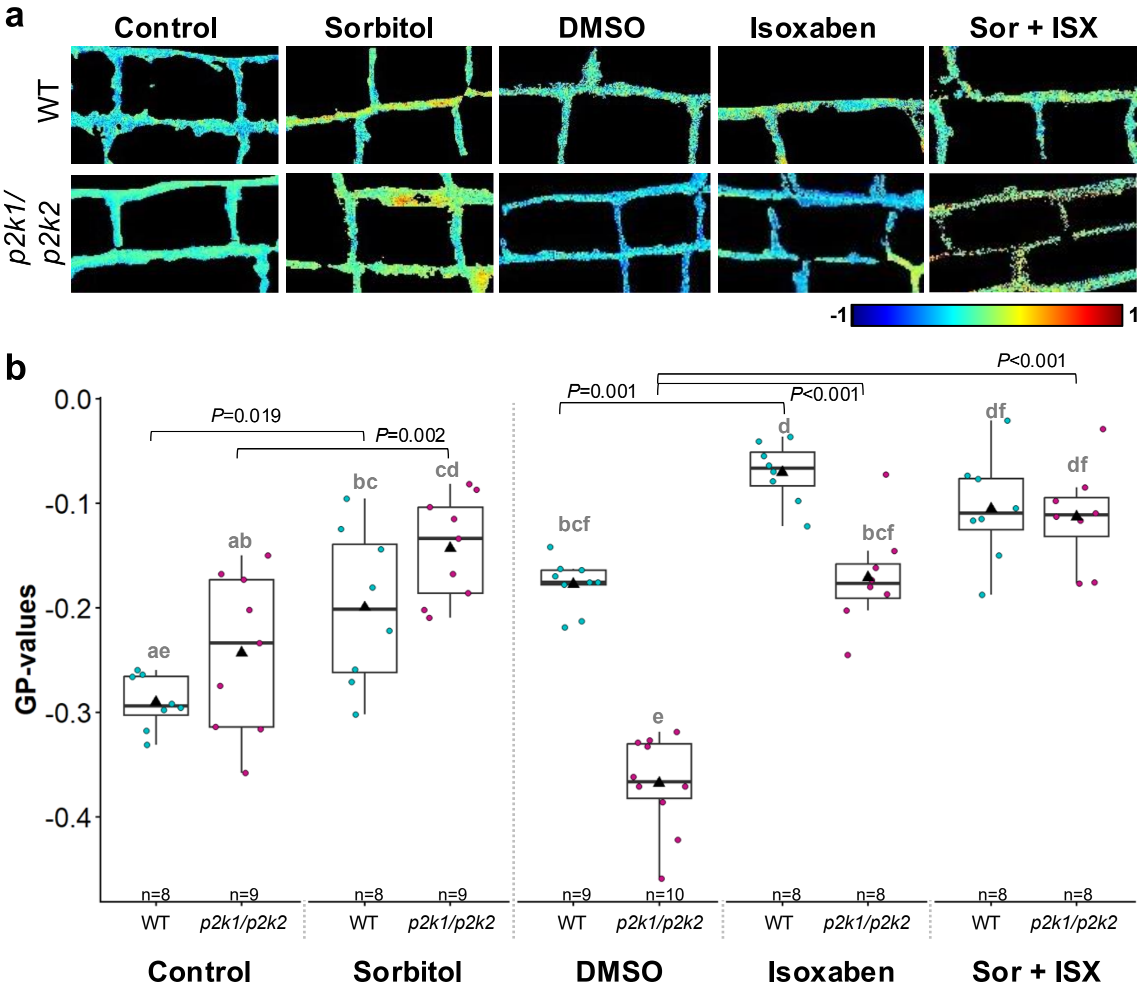


**Supplementary Figure 9. Plasma membrane polarity pattern in cortex cells of Col-0 and *p2k1/p2k2* root elongation zones**. **a,** Graphical representations of wildtype and *p2k1/p2k2* cortex cells (digitally magnified from Supplementary Fig. 8b) stained with 40μM C-Laurdan after control, mock (DMSO), 0.3M sorbitol, 0.6μM ISX, and combined treatments for four hours. **b,** GP values of plasma membrane of wild-type and *p2k1/p2k2* cortex cells. Each data point indicates mean of cortex cell GP-values per individual root. Error bars = standard deviation. Statistics: one-way ANOVA with Post Hoc – Tukey HSD (*P*-values indicate the statistical significance of differences between treated roots and their respective control or DMSO group, *alphabet* refers to significance between all samples).
